# Common Mechanism of SARS-CoV and SARS-CoV-2 Pathogenesis across Species

**DOI:** 10.1101/2021.05.14.444205

**Authors:** Alexandra Schäfer, Lisa E. Gralinski, Sarah R. Leist, Emma S. Winkler, Brea K. Hampton, Michael A. Mooney, Kara L. Jensen, Rachel L. Graham, Sudhakar Agnihothram, Sophia Jeng, Steven Chamberlin, Timothy A. Bell, D. Trevor Scobey, Laura A. VanBlargan, Larissa B. Thackray, Pablo Hock, Darla R. Miller, Ginger D. Shaw, Fernando Pardo Manuel de Villena, Shannon K. McWeeney, Stephanie A. Montgomery, Michael S. Diamond, Mark T. Heise, Vineet D. Menachery, Martin T. Ferris, Ralph S. Baric

**Author notes:** Co-first authors (Alexandra Schäfer, Lisa E. Gralinski, Sarah R. Leist). Co-senior authors.

## Abstract

Sarbecovirus (CoV) infections, including Severe Acute Respiratory CoV (SARS-CoV) and SARS-CoV-2, are considerable human threats. Human GWAS studies have recently identified loci associated with variation in SARS-CoV-2 susceptibility. However, genetically tractable models that reproduce human CoV disease outcomes are needed to mechanistically evaluate genetic determinants of CoV susceptibility. We used the Collaborative Cross (CC) and human GWAS datasets to elucidate host susceptibility loci that regulate CoV infections and to identify host quantitative trait loci that modulate severe CoV and pan-CoV disease outcomes including a major disease regulating loci including *CCR9. CCR9* ablation resulted in enhanced titer, weight loss, respiratory dysfunction, mortality, and inflammation, providing mechanistic support in mitigating protection from severe SARS-CoV-2 pathogenesis across species. This study represents a comprehensive analysis of susceptibility loci for an entire genus of human pathogens conducted, identifies a large collection of susceptibility loci and candidate genes that regulate multiple aspects type-specific and cross-CoV pathogenesis, and also validates the paradigm of using the CC platform to identify common cross-species susceptibility loci and genes for newly emerging and pre-epidemic viruses.

## Main Text

Natural host genetic variation regulates disease severity following most viral infections, yet the specific genes and loci that regulate differential disease outcomes are largely unknown across susceptible species^1,2^. Coronaviruses (CoVs) are significant human and animal pathogens; six CoVs (three human, three swine) have emerged or expanded their geographic range in the 21^st^ century^3,4^. The most impactful emergent human CoVs (SARS-CoV and SARS-CoV-2) are group 2B coronaviruses, which likely emerged from bats to cause worldwide human epidemic or pandemic respiratory infections, leading to substantial morbidity and mortality^5,6^. Moreover, many high-risk group 2B Sarbecoviruses (SARS-like viruses) and group 2C MERS-like bat CoVs are poised for future human emergence events^7–9^. The Sarbecovirus subgenus is clustered into four clades that include the clade I SARS-CoV and high-risk SARS-like Bat CoVs (BtCoV), clade II SARS-like BtCoVs like HKU3, and clade III SARS-CoV-2 (**Figure 1a**)^10^. Sarbecoviruses vary widely in their ability to cause human and animal disease^11^. SARS-CoV caused ~8,000 infections with a 10% mortality rate, while SARS-CoV-2 has infected ~132 million, leading to ~2.9 million deaths to date^12,13^. SARS-CoV-2 infections cause asymptomatic to life-threatening disease outcomes, supporting a role for inter-host genetic control of emerging viral disease outcomes in both humans and mice^14–16^. Thus, understanding the functions of natural host variants in genes that regulate susceptibility and disease severity after diverse Sarbecovirus infections may reveal common genetic loci that regulate wildtype and variant virus pathogenic outcomes across species, inform threat potential, and reveal novel targets for therapeutic intervention.

**Figure 1.**
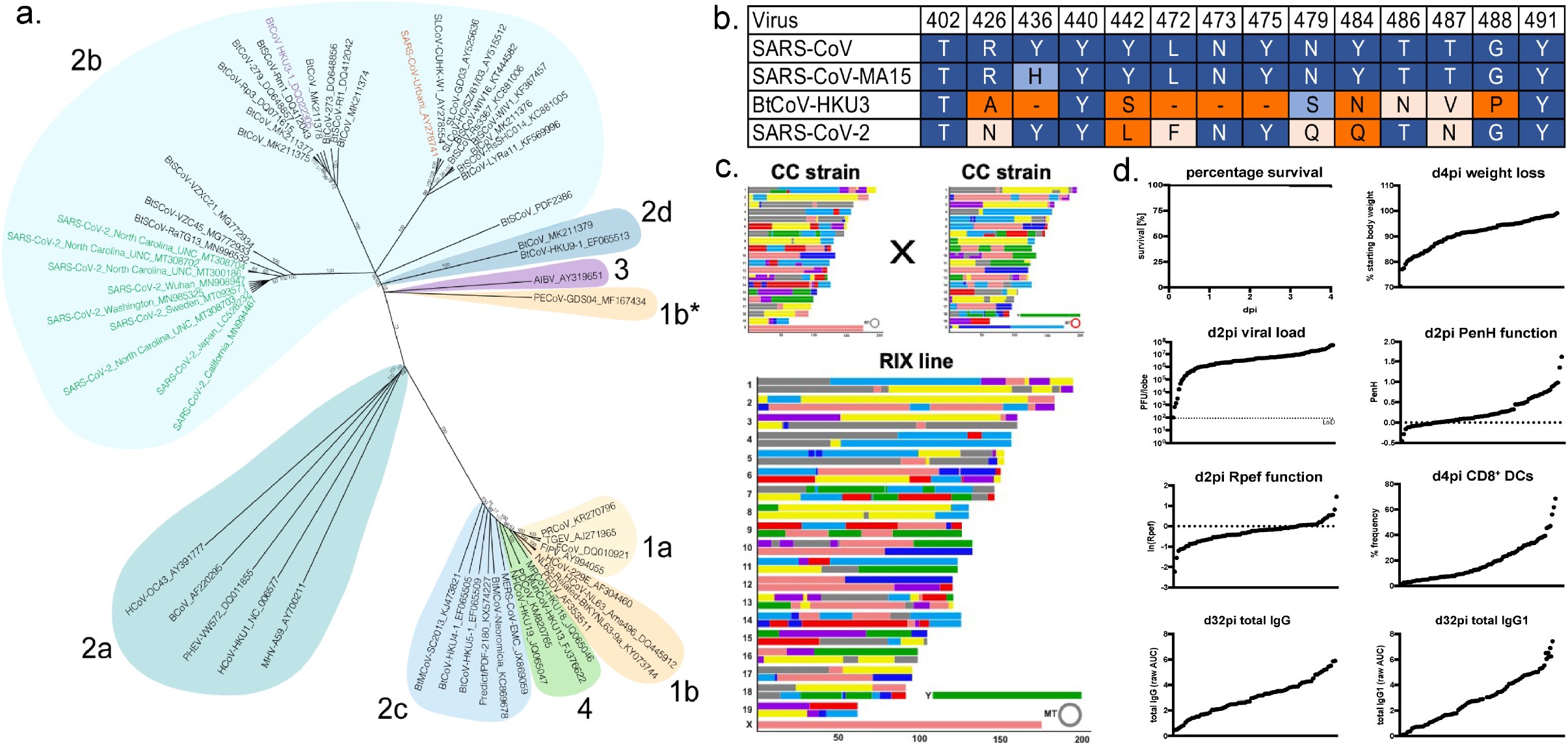
Coronavirus and host-genetic model systems: a-b. Spike phylogeny of representative coronaviruses and RBD alignments SARS-CoV, HKU3, and SARS-CoV-2 with emphasize on ACE2 binding residues; c.-d. The RIX Collaborative Cross Screen: The design of CC-RIX panel and the phenotypic distribution of disease phenotypes after SARS-MA and HKU3-MA infection in the CC-RIX panel. **a.** The Spike protein sequences of selected coronaviruses were aligned and phylogenetically compared. Coronavirus genera are grouped by classic subgroup designations (1a-b, 2a-d, 3, and 4). PECoV is designated as 1b* because of its distinctive grouping compared with more conserved proteins. Branches in each tree are labeled with consensus support values (in %). Sequences were aligned using free end gaps with the Blosum62 cost matrix, and the tree was constructed using the neighbor-joining method based on the multiple sequence alignment in Geneious Prime. Numbers following the underscores in each sequence correspond to the GenBank Accession number. **b.** Spike receptor-binding domain (RBD) alignments were performed in Geneious using free end gaps with the Blosum62 cost matrix, and 14 ACE2-critical interacting residues are highlighted in the chart. Residues in the chart are color-coded based on conservation to the SARS-CoV residue (conservation based on Blosum62). **c.** The Collaborative Cross is an octo-parental genetic reference panel. The population captures ~40 million SNPs and small InDels without blind spots across the genome. Each individual strain’s unique combination of haplotypes across the genome results from the independent recombinations each strain experienced in their generation. As such, each CC strain represents a unique combination of Coronavirus susceptibility alleles. **d.** Phenotype distribution in CC-RIX panel during SARS-MA infection. Each dot represents the mean of individual CC-RIX strains.

The angiotensin-converting enzyme 2 (ACE2) receptor interacts with the Spike protein (S) receptor binding domains of SARS-CoV, SARS-CoV-2 and many BtCoV Sarbecoviruses, but not the clade II HKU3 strain (**Figure 1b**)^17^ As many of these strains do not produce disease in mice, reverse genetics and serial passaging were used to select for SARS-CoV-MA15 (SARS-MA), HKU3-MA, and SARS-CoV-2 MA10 strains that replicate efficiently and produce severe disease in mice^11,18,19,20^. Mouse genetic reference populations (GRPs) have been employed as highly relevant models of human disease, and coupled with systems genetics data algorithms, to identify host susceptibility loci, genes, genetic networks and higher-level genetic interactions that regulate phenotypic variation and disease severity^21–23^. Among these mouse GRPs, the Collaborative Cross (CC) genetic reference population encodes over 44 million single nucleotide polymorphisms (SNPs), 4 million insertions and deletions (InDels), as well as several thousand novel variants (both SNPs as well as InDels) present in single strains^24,25^. Building from our prior work, genetically mapping quantitative trait loci in incipient mice from CC strains (Pre-CC^26^, **Table 1**), we designed a panel of 115 F1 crosses between different CC strains (CC-RIX; an outbred population, like humans, but reproducible (**Figure 1c**)) to identify loci controlling Sarbecovirus pathogenesis and adaptive immune responses. We used our mouse-adapted CoV models, including SARS-CoV MA15 (SARS-MA), HKU3-MA, and SARS-CoV-2 MA10, which replicate efficiently and produce severe disease in mice^11,18,19,20^, to overcome the host-specific angiotensin-converting enzyme 2 (ACE-2) interaction with CoV Spike protein.

**Table1.** List of significant QTLs

| QTL ID | Population | Chromosome [Mb] | Phenotype(s) (days post infection- dpi) | Haplotype(s)<sup>+</sup> | phenotypic variation [%] |
| --- | --- | --- | --- | --- | --- |
| **Pre-CC** |  |  |  |  |  |
| *HrS1* | Pre-CC | chr3:18.3-26.8 | Perivascular cuffing (4dpi) | BH/ACDEFG | 26% |
| *HrS2* | Pre-CC | chr16:31.5-36.7 | Viral titer (4dpi) | G/ABCDEFH | 22% |
| *HrS3* | Pre-CC | chr15:72.2-76.0 | Eosinophilia (4dpi) | A/BCDEFGH | 26% |
| *HrS4* | Pre-CC | chr13:52.7-54.9 | Perivascular cuffing (4dpi) | F/ABCDEGH | 21% |
| **CC-RIX** |  |  |  |  |  |
| *HrS10* | CC-RIX | chr13: 20.3- 23.7 | HKU3-MA mortality | FG/ABCDEH | OR* 6.13 |
| *HrS11* | CC-RIX | chr5: 112.9-118.8 | HKU3-MA mortality | C/ABDEFGH | OR* 10.45 |
| *HrS12* | CC-RIX | chr15: 51.5- 75.6 | HKU3-MA weight loss (4dpi) | G/ABCDEFH | 3.4% |
| *HrS13* | CC-RIX | chr16:18-51.1 | SARS viral titer (2dpi) | C/G/ABDEFH | 15.94% |
| *HrS14* | CC-RIX | chr11: 9.05- 28.66 | PenH (2dpi) | D/CEH/ABFG | 11.65% |
| *HrS15* | CC-RIX | chr11: 3.26- 28 | Rpef (4dpi) | ABCEFGH/D | 6.48% |
| *HrS17* | CC-RIX | chr17: 27.4- 78.3<br>chr17: 27.4- 78.3 | IgG1 (N) (32dpi)<br>Total IgG (N) (32dpi) | A/BCDEFH/G | 19.97%<br>18.53% |
| *HrS18* | CC-RIX | chr16: 18.1- 43.8 | Total IgG (N) (32dpi) | ABCDEFH/G | 22.75% |
| *HrS19* | CC-RIX | chr10: 17.4- 130.5 | Total IgG (N) (32dpi) | ABCDEH/FG | 10.71% |
| *HrS20* | CC-RIX | chr3: 3.1- 35.4 | Total IgG (N) (32dpi) | G/ABCDEFH | 10.83% |
| *HrS22* | CC-RIX | chr6:67.2- 85 | CD8<sup>+</sup> DCs [%] (4dpi) | F/ABCDEGH | 8.17% |
| **CC011xCC074-F2** |  |  |  |  |  |
| *HrS24* | CC011xCC074-F2 | chr4:32.95-114.54 | Mortality | CC011: G,H,C<br>CC074:A/H,<br>A/D, D, E, F | OR* 4.34 |
| *HrS25* | CC011xCC074-F2 | chr4: 6.38-17.97 | Weight loss in males (4dpi) | CC011: G, B<br>CC074: A | 12.57% |
| *HrS26* | CC011xCC074-F2 | chr9:74.94-124.06<br>chr9:117.38-124.07<br>chr9:116.24-124.07 | Mortality<br>Hemorrhage (4dpi)<br>PenH (2dpi) | CC011: B, G<br>CC074: A | OR* 3.15<br>10.24%<br>7.76%<br>11.8% |
|  |  | chr9:111.54-122.63 | Periph. neutrophils (4dpi) |  | 12.39% |
|  |  | chr9:111.54-122.63 | Periph. lymphocytes (4dpi) |  |  |
| *HrS27* | CC011xCC074-F2 | chr11:26.44-80.76<br>chr11:17.89-80.76 | Periph. neutrophils (4dpi)<br>periph. lymphocytes (4dpi) | CC011: H, B, H<br>CC074: C | 5.5%<br>5.6% |
| *HrS28* | CC011xCC074-F2 | chr15:58.66-74.04 | PenH (2dpi) | CC011: H<br>CC074: B | 8.48% |
| **CC003xCC053-F2** |  |  |  |  |  |
| *HrS5* | CC003xCC053-F2 | chr18:24.76-51.25<br>chr18:31.63-51.25<br>chr18:42.85-73.43<br>chr18:31.63-65.17<br>chr18:42.85-65.17<br>chr18:24.76-51.2<br>chr18:24.76-51.2 | Weight loss (3dpi)<br>Weight loss (4dpi)<br>Hemorrhage (4dpi)<br>Perivascular cuffing (4dpi)<br>Edema (4dpi)<br>Viral titer (4dpi) | CC003: A, G<br>CC053: C/G, G, B, B/D, B/C, D | 6%-13.2% |
| *HrS6* | CC003xCC053-F2 | chr9:121.77-end | Weight loss (3dpi) | CC003: B, B/D<br>CC053: H | 7% |
| *HrS7* | CC003xCC053-F2 | chr7:83.45-129.93<br>chr7:69.79-96.66 | Weight loss (4dpi)<br>Viral titer (4dpi) | CC003: B, C, B/C<br>CC053: G/H, H, D | 6.8%<br>14.4% |
| *HrS8* | CC003xCC053-F2 | chr12:88.54-103.21 | Viral titer (4dpi) | CC003: D<br>CC053: F/H | 4.27% |
| *HrS9* | CC003xCC053-F2 | chr15:10.7-57.3 | Hemorrhage (4dpi) | CC003: G, E/G, G<br>CC053: D, C, F | 9.4% |
| *HrS23* | CC003xCC053-F2 | chr4:35.61-104.04<br>chr4:35.61-104.04 | Hemorrhage (4dpi)<br>Weigh loss (4dpi) | CC003: H, E, G, H<br>CC053: F, G, F | 6.86%<br>6.22% |

\*QTL influencing mortality calculate an Odds Ratio (OR) for disease rather than a % of variation explained. For both the CC-RIX and the CC-F2, Odds Ratios for specific loci are calculated with a full model of all mortality loci, to better estimate their independent effects.
\*Haplotype effects are described for each QTL. For the Pre-CC and CC-RIX, the haplotypes are separated based on the allele effect splits between founder haplotypes (A=A/J, B=C57BL/6J, C=129S1/SvImJ, D=NOD/ShiLtJ, E=NZO/HiLtJ, F=CAST/EiJ, G=PWK/PhJ, H=WSB/EiJ). In CC-F2 crosses, the haplotypes listed are those present in the given parent strains from the proximal to distal ends of each region

## Results

### Phenotypic distributions, genomics scans, and allele effects maps for 3 traits across the CC-RIX

We infected the CC-RIX population with two genetically distinct Sarbecoviruses, which included the clade I SARS-MA and clade II HKU3-MA strains, respectively (**Figure 1a-b**)^11,27^. Groups of CC-RIX mice were inoculated with 5×10^3^ plaque-forming units (PFU) of SARS-MA, and viral burden, clinical disease (*e.g*., weight loss, mortality, and respiratory function), antibody titers, and immune cell infiltrates were measured at multiple timepoints post-infection (ranging from 2-32 days) (**Figure 1d, Figure 2**). A matched cohort of CC-RIX was inoculated with 1×10^5^ PFU of HKU3-MA and evaluated for mortality and weight loss through day 4 post infection (**Figure 2a**). In both studies, virus challenge elicited an array of disease phenotypes, ranging from clinically inapparent infection to lethal outcomes within the first 4 days of infection. We estimated genetic contributions for many of these traits and determined that heritability for these responses was 44.4%-80.9%, estimates that agree with previous CC studies^26,28–31^. Importantly, the various CC-RIX disease phenotypes measured in response to infection appeared relatively uncorrelated (**Figure 2a-c**), suggesting that there are many independent genetic factors driving these responses. We next conducted genetic mapping to identify both genomic regions and specific founder haplotypes driving various aspects of SARS-MA and HKU3-MA disease responses. Given the likely complex genetic architecture underlying these phenotypes, we identified those loci surpassing community-accepted significance thresholds (p<0.33), with distinct allele effects (**Figure 2a-c, Table 1**)^32^. We identified 11 distinct and high-confidence loci in our RIX population affecting weight loss, mortality, titer, antibody responses or respiratory function after SARS-MA infection or weight loss and mortality following HKU3-MA infection. The effect sizes of these loci varied from 3-23% of the total trait variation (that is, largely moderate effect sizes), the loci were located in different genomic regions with different causal founder alleles, and most loci primarily impacted one or a few traits in this population (**Table 1**). The number of independent loci and their trait-specific impacts are consistent with our earlier observations of little to no correlation between disease states across the RIX lines (**Figure 2a-c**). Together, this analysis highlights the genetic complexity driving CoV disease outcomes and immunity and also the inability of any single animal model of CoV disease to fully address all aspects of the disease response.

**Figure 2.**
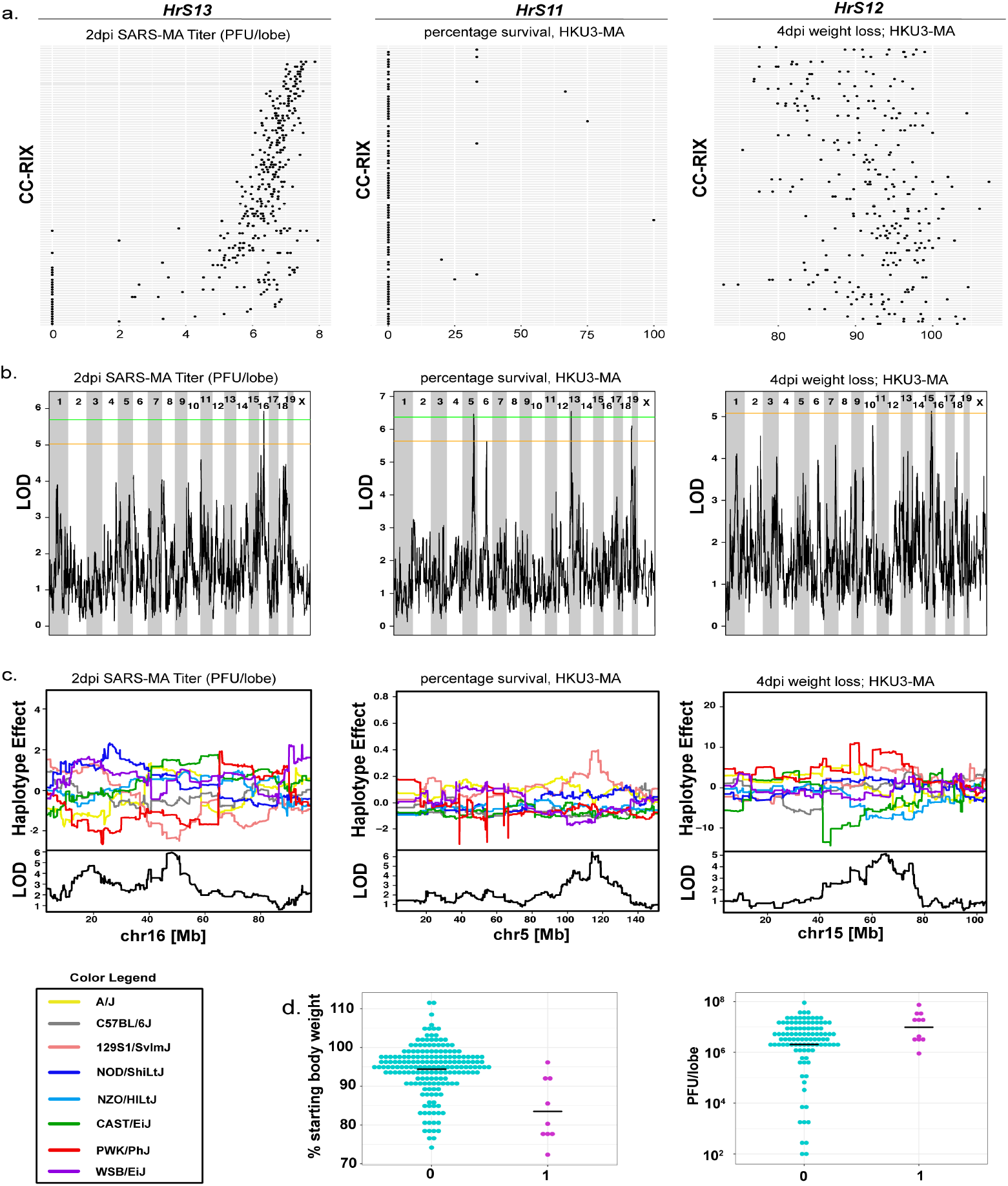
Phenotypic distributions, genomics scans, and allele effects maps for 3 traits across the CC-RIX. **a.** *HrS13*, SARS-CoV lung titer at 2 dpi, *HrS11*, percentage survival following HKU3-MA infection, and *HrS12*, weight loss at 4 dpi with HKU3-MA. For all 3 panels, CC-RIX strains are sorted by ascending 2 dpi SARS-MA lung titers, showing the general lack of correlation between coronavirus disease responses. **b.** Genome scans showing the LOD traces, as well as significance thresholds (p=0.33 (orange) and p=0.2 (green)) for the same traits listed above. We identified *HrS13* (ch16) for SARS-CoV titer, *HrS11* (chr5) and *HrS10* (chr13) for HKU3-MA mortality, and *HrS12* for HKU3-MA weight loss. **c.** Allele effects at each of these loci showing causal haplotypes for *HrS13* (where 129S1/SvImJ (pink) and PWK/PhJ (red) alleles cause a reduced titer, *HrS10* where CAST/EiJ (green) and PWK/PhJ (red) alleles cause increased mortality, *HrS11* where a 129S1/SvImJ (pink) allele causes increased mortality, and *HrS12* where a PWK/PhJ (red) allele causes decreased weight loss. **d**. We identified relationships between *HrS10* and SARS-related weight loss and clinical disease (shown is weight loss at 5 dpi), as well as *HrS11* and SARS-MA titer (shown here is titer at 2 dpi), 0 = low response haplotype, 1 = 1 copy of the high response/PWK haplotype. Each dot represents data from an individual animal.

To investigate the possibility of pan-sarbecovirus susceptibility loci, we evaluated whether any of our SARS-MA phenotypes also were associated with the haplotypes driving HKU3-MA mortality (**Figure 2d**, **Table 1**). For these analyses, we binned CC-RIX lines based on their diplotypes at these HKU3-MA diseases associated loci, then determined whether these diplotype bins provided an improved fit to the relationship between SARS-MA phenotypes and the CC-RIX IDs themselves. For example, was there any genetic signal at quantitative trait loci (QTL) *HrS10* or *HrS11* that was associated with differential SARS-MA disease when simplifying the underlying causal model; a method we have used to find additional QTLs with small effect sizes in genetically complex populations^29^. *HrS10* was associated with enhanced SARS-MA weight loss, disease and mortality at day 2 post-infection (2 dpi), whereas *HrS11* had moderate associations with SARS-MA lung titers at both 2 and 4 dpi (**Figure 2d**). These results indicate that common susceptibility loci regulate disease severity across two genetically distinct Sarbecovirus clades, and that *HrS10* and *HrS11* are both associated with HKU3-MA-induced mortality, but likely contribute to virus-induced disease through different mechanisms.

### Disease phenotypes in parental strains CC011 and CC074 and identification of a quantitative multitrait locus on chromosome 9 in the CC011xCC074-F2

Concurrent with our large CC-RIX screen, we identified a pair of inbred CC strains showing highly divergent susceptibilities to SARS-CoV: the disease resistant CC011/Unc and the highly susceptible CC074/Unc strain (CC011 and CC074 from here on, respectively). After challenge with 1×10^4^ PFU SARS-MA, these strains exhibited marked differences in clinical and virological disease phenotypes (*e.g*., hemorrhage, weight loss, virus titer, mortality, circulating immune cells), and all CC074 mice developed lethal disease by 4 dpi post infection (**Figure 3a, left panel**). Relevant for the current pandemic, these parental strains showed similar severe infection phenotypes during SARS-CoV-2 MA10 infection (**Figure 3a, right panel, Figure S1**). We generated 403 F2 mice by intercrossing these strains (**Figure S2**) and inoculated them intranasally at 9-12 weeks with 1×10^4^ PFU of SARS-MA. These F2 mice showed an expanded range of disease responses relative to their parent CC strains, including mortality, weight loss, titer, respiratory function, circulating immune cell and hemorrhage phenotypes (**Figure S3, Figure 3b**). We conducted QTL mapping in these F2 mice and identified five significant QTL segregating in this population (*HrS24-28*, **Table 1**), and supply information on other potential loci (p<0.33, **Table S1**). Most of these loci impacted multiple aspects of the SARS-MA disease response during this infection time course (for example analysis of *HrS26* on chromosome 9 indicated the locus contributed to mortality at 4 dpi (Odds ratio (OR) of 3.15), as well as lung hemorrhage or congestion (10.2% of population variation), airflow restriction at 2 dpi (7.8% PenH), as well as peripheral neutrophil (11.8%) and lymphocyte (12.4%) levels) at 4 dpi (**Figure 3c, Table 1**). Recently, three Genome-wide Association Study (GWAS) in humans identified a locus associated with respiratory failure. This locus (encompassing genes such as *SLC6A20, LZTFL1, FYCO1, CXCR6, XCR1*, and *CCR9*) is syntenous with a more proximal region of our chromosome 9 locus^33–35^. *CCR9* emerged as a strong candidate based on the integration of our data with these studies and the presence of nonsynonomous SNPs in *CCR9* as well as synonymous mutations in regulatory flanking sequences.

**Figure 3.**
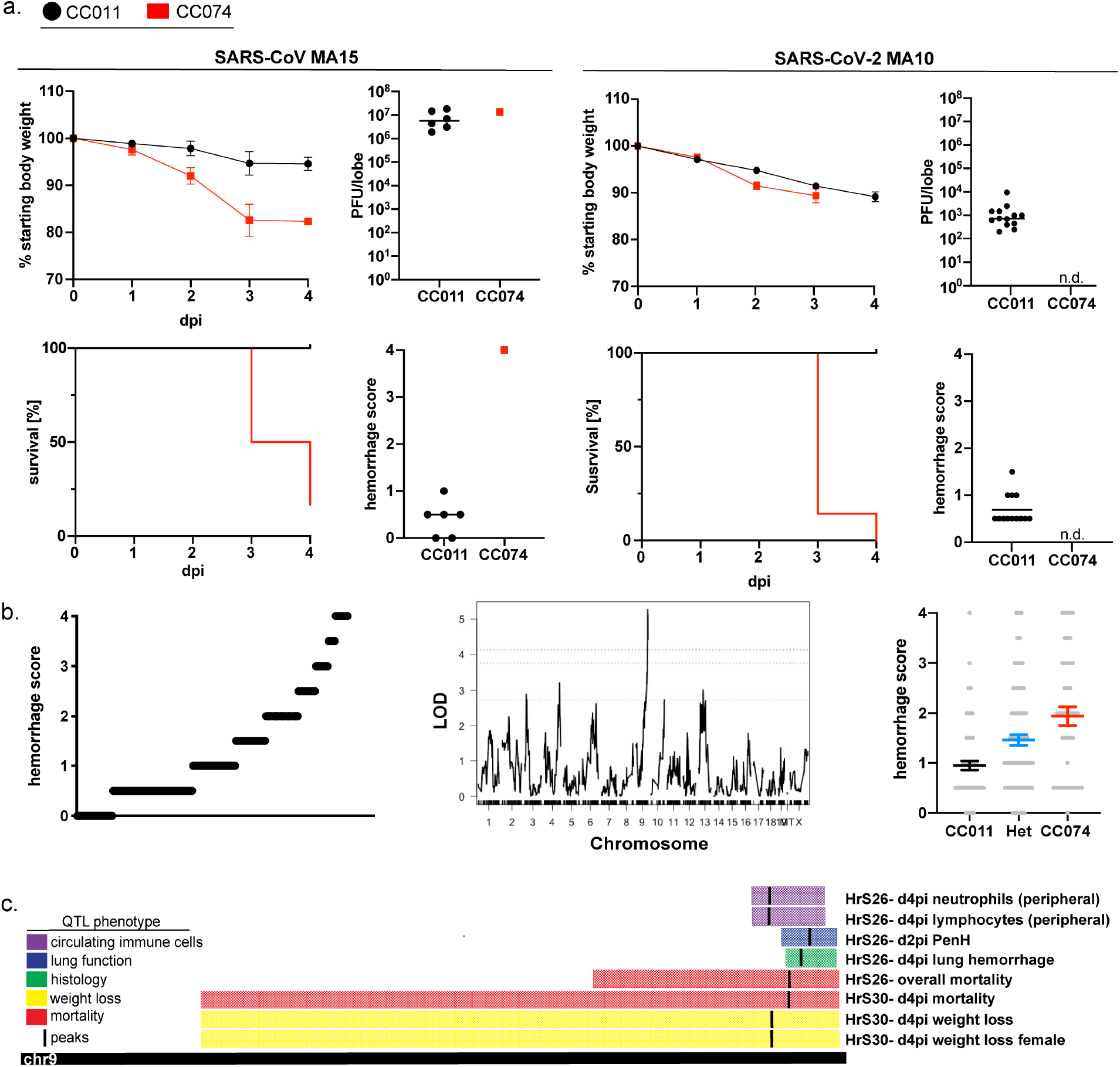
Disease phenotypes in parental strains CC011 and CC074 and identification of a quantitative multitrait locus on chromosome 9 in the CC011xCC074-F2. **a.** CC011 and CC074 were identified as parental CC strains for a F2-screen based on their contrasting SARS-MA induced phenotypes. Shown are weight loss, lung viral titer, percentage survival, and lung hemorrhaging following SARS-MA infection (n=6 for CC011 and CC074, respectively) and SARS-CoV-2 MA10 (n=13 for CC011 and n=13 for CC074). **b.** Shown as example is the phenotypic distribution for lung hemorrhage following SARS-MA infection in the CC011xCC074-F2 panel. The genomic scan shows the LOD traces and significance thresholds (p=0.95 (black), p=0.90 (blue), and p=0.50 (green)). The allele effect of the lung hemorrhage phenotype is broken out based on the homozygous CC011 genotype, the homozygous CC074 genotype, or the heterozygous genotype (n=234 individual CC011xCC074-F2 mice). **c.** A quantitative multitrait locus with major effect was identified on chr9 (74.9–124 Mb), which affected mortality, weight loss, hemorrhage, lung function, and peripheral hematology. CC011/Unc has a C57BL/6/PWK haplotype and CC074/Unc has an A/J haplotype in this QTL region.

### Identification of *CCR9* as a major susceptibility allele during SARS-CoV-2 infection

To better understand how our contrasting CC strains and this locus regulates SARS-CoV-2 disease, we inoculated CC011, CC074, C57BL/6NJ and *CCR9^-/-^* mice on a C57BL/6NJ background with SARS-CoV-2 MA10^20^. CC011 and CC074 mice infected with SARS-CoV-2 MA10 showed concordance in their SARS-MA disease response phenotypes, including lethality in the CC074 susceptible line. The *CCR9^-/-^* mice developed more severe clinical disease (**Figure 4a, Figure S4a**), exhibited increased virus titers (**Figure 4b**), weight loss, mortality and prolonged pulmonary dysfunction and severe lung pathology as measured by whole body plethysmography (**Figure 4c**), lung hemorrhage (**Figure S4b**), and lung damage (**Figure S4c, d and Figure S4e**) relative to their wild-type controls, supporting its important role in protection from disease. Analysis of the cytokine profile in lungs and serum by multiplex immune-assay showed increased subsets of cytokines and chemokines that are involved in promoting allergic airway inflammation, including IL9, IL13, Il17, CCL2, CCL3, CCL5, G-CSF, and Eotaxin either in the lung tissue, serum, or both (**Figure 4f and Figure S5b**). By flow cytometric analysis, the composition of infiltrating cells into the lung tissue (**Figure 4e**) and bronchoalveolar (BAL) fluid (**Figure S5a**) of *CCR9^-/-^* mice showed a significant increase of CD4^+^ T cells, CD8^+^ effector T cells, inflammation-promoting CD11^+^ DCs and eosinophils at 6 dpi, consistent with an allergic airway inflammatory response. Although originally found to play an important role in chronic gut inflammation, *CCR9* is mainly expressed on lymphocytes, dendritic cells (DCs) and monocytes/macrophages^36,37^, and also participates in early allergic airway inflammation including the migration and proliferation of eosinophils and lymphocytes^38^. In addition *CCR9^+^* DCs are implicated in regulating inflammation, alloimmunity, and autoimmunity^36^. *CCR9^-/-^* mice develop chronic inflammatory responses and CD11b^+^ inflammatory macrophages contribute to the pathogenesis of liver fibrosis via the *CCR9/CCL25* axis^39,40^. As inflammatory macrophages contribute significantly to increased SARS-CoV pathogenesis in mice^41^, together, these data target the *CCR9/CCL25* axis as a major driver of SARS-CoV-2 pathogenesis across species and validate *CCR9* as a driver of the human Chr3 susceptibility loci and mouse Chr9 susceptibility locus.

**Figure 4.**
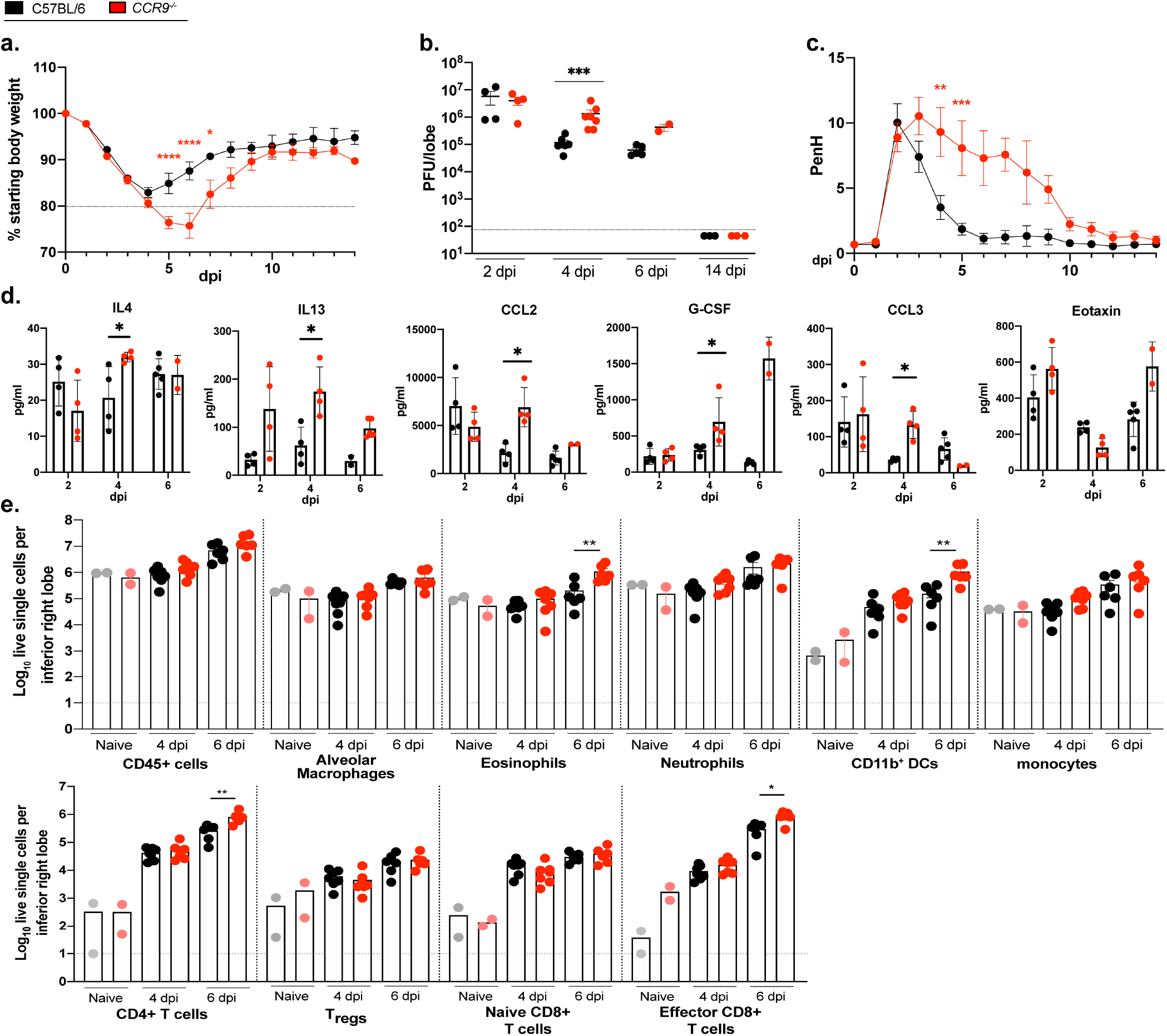
Identification of *CCR9* as a major susceptibility allele during SARS-CoV-2 infection. Infection of *CCR9^-/-^* mice with SARS-CoV-2 MA10 showed significant differences in weight loss (**a.**), viral burden in the lung (**b.**), lung function (**c.**), and cytokine/chemokine distribution (**d.**) in the as well as in the compositions of lung infiltrating immune cells (**e.**) (n=19 *CCR9^-/-^* and n=19 C57BL/6NJ; with n=4 for 2 dpi, n=7 for 4 dpi, n=5 for 6 dpi, and n=3 for 14 dpi for weight loss, viral burden, congestion score, multiplex immune-assay and lung pathology; n=8 for lung function testing; n=12 CCR9^-/-^ and n=13 C57BL/6NJ with n=2 each for mock, n=5-6 for 4 dpi, and n=5 for 6 dpi for analysis of infiltrating cells). Data were analyzed using two-way-ANOVA (weight loss, lung function) and Mann-Whitney test (titer, congestion score, pathology scores, FACS analysis, and cytokine/chemokine analysis; *p<0.05, **p<0.01.

A key goal in animal studies is to identify relevant models of human disease. Syntenic genome regions between humans and rodents often regulate a number of infectious and chronic diseases, and our analysis of *HrS26* extends this important pattern^42–44^ to sarbecovirus infections. All told, the synteny between human Chr3 and mouse Chr9, the effect sizes of the loci identified in this F2 (between 5-15% of the population-wide phenotypic variation in this F2, **Table 1**), as well as the prevalence of loci impacting multiple aspects of SARS-MA associated disease, highlight how sorting of multiple susceptibility alleles into individual CC strains model unique aspects of the genetic control of disease responses. Furthermore, the presence of alleles of at least six of the CC founder strains segregating across these loci (and often in opposite directions: a C57BL/6J allele is protective at *HrS26* and *HrS27* but exacerbating at *HrS28*, **Table 1**) highlights the utility of using genetically complex but reproducible models of disease.

### Identification of major effect locus on chromosome 4 and of *Trim14* as a susceptibility gene during SARS-MA and SARS-CoV-2 MA10 infection

Next, we revisited a previous CC-F2 intercross, CC003/UncxCC053/Unc (named CC003 and CC053 from here on) conducted by our group^45^, and utilized our refined analysis pipelines once the original SARS-MA disease loci (*HrS5-9*) were statistically accounted for. This re-analysis allowed us to identify an additional locus (*HrS23* on chr4) impacting both weight loss and hemorrhagic damage to the lungs as determined by gross pathology (**Figure 5a, b**), as well as several other suggestive QTLs (**Table S1**). The chr4 locus also overlapped with the mortality QTL *HrS24* identified in CC011xCC074-F2 cross (**Figure 5c**). SNP variation between CC003 and CC053, as well as between CC011 and CC074 in this locus pointed to *Trim14* as a likely candidate gene driving these differences in SARS-CoV disease. Previous work identified *Trim14* as a key docking platform in the context of MAVS signaling^46,47^. We used CRISPR/Cas9 targeting to edit *Trim14* in C57BL/6J mice, create a functional knockout (**Figure S6**), and evaluate its role following infection (**Figure 5d, top panel**). *Trim14^Δ47/Δ47^* mice inoculated with 1×10^5^ PFU of SARS-MA had a modest increase in pathogenesis relative to C57BL/6J control mice. At 3 and 4 dpi, *Trim14*-deficient mice had increased weight loss, which corresponded with increases in viral titer within the lung at 2 and 4 dpi. This result show that an absence of *Trim14* affects viral clearance. Similarly, *Trim14^Δ47/Δ47^* mice inoculated with SARS-CoV-2 MA10 also sustained modest increases in weight loss and a delayed recovery phenotype when compared to C57BL/6 mice (**Figure 5d, bottom panel**). However, the difference in viral titer seen at early times post SARS-MA infection was not observed with SARS-CoV-2 MA10. Together, these data suggest that Trim14 has a shared role in attenuating Sarbecovirus disease potential, but that this effect varies between viruses. Taken together, the data demonstrates the potential of identifying common and shared QTLs among group 2B coronaviruses.

**Figure 5.**
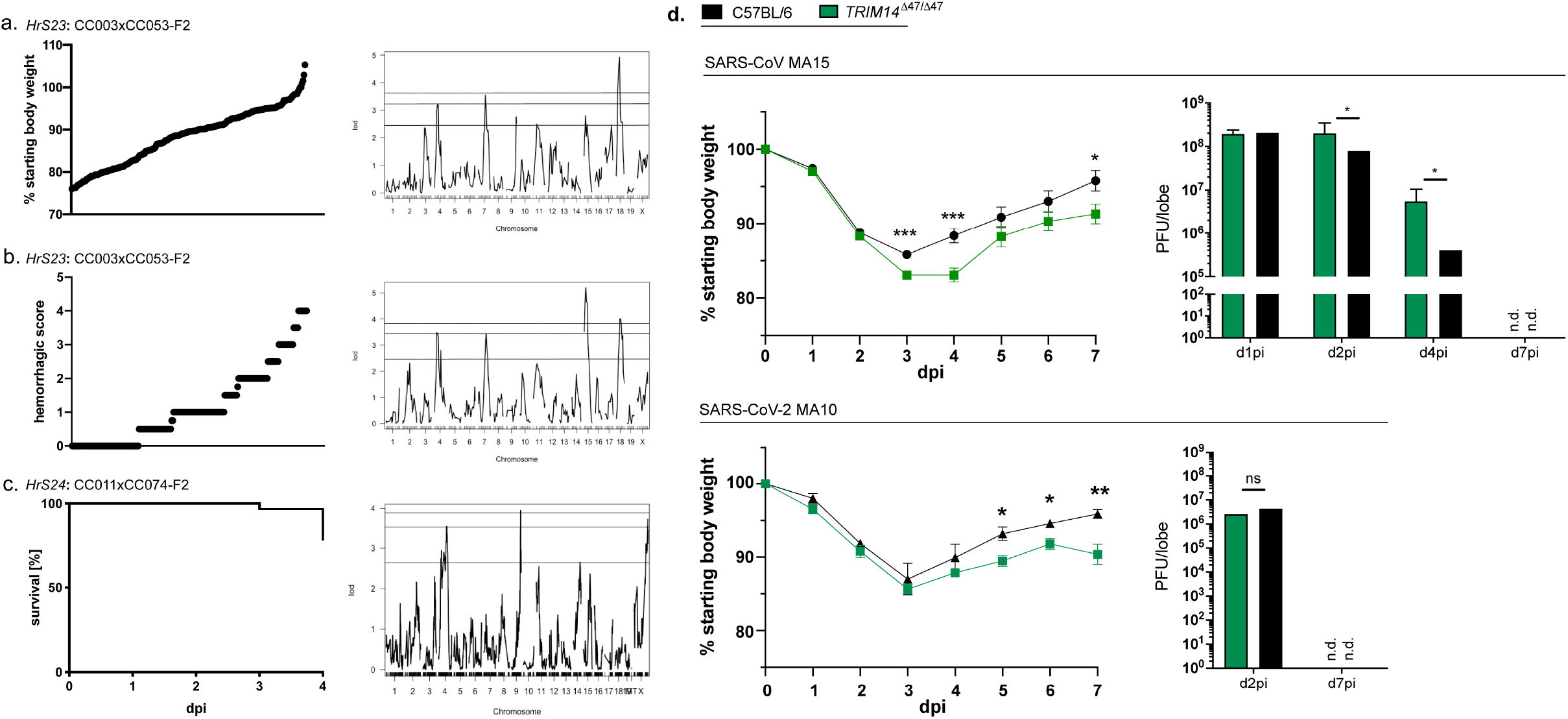
Identification of major effect locus on chromosome 4 and of *Trim14* as a susceptibility gene during SARS-MA and SARS-CoV-2 MA10 infection. **a.** Phenotypic distribution and genomic scan for *HrS23* (4dpi weight loss in CC003xCC053-F2, **b.** *HrS23* (4dpi lung hemorrhage in CC03xCC053-F2), and **c.** *HrS24* (overall mortality in CC011xCC074-F2. **d.** Infection of *Trim14^Δ47/Δ47^* mice with SARS-MA showed significantly more weight loss and an increase in viral load in the lung compared to C57BL/6J mice over the course of a 7-day infection. A similar trend of infection progression was observed in *Trim14^Δ47/Δ47^* mice infected with SARS-CoV-2 MA10; n=10 *Trim14^Δ47/Δ47^* and n=10 C57BL/6J for weight loss and viral load studies with SARS-CoV-2 MA10 studies; data was analyzed via student t test, *p<0.05, **p<0.01, ***p<0.005).

## Discussion

Across these studies, we describe several dozen loci impacting different disease responses, including several which show broad responses to all tested Sarbecoviruses. We demonstrate connections to human GWAS studies, and as such these data represent a resource for future comparative studies of Sarbecovirus pathogenesis between humans and animals. Our study highlights the power of using animal GRPs to understand the role of host genetic variation on infectious diseases, generate new models of differential disease, probe the role of individual genes in disease progression, and provide mechanistic insight into the role of specific host genes and viral strains in regulating pathogenesis across species. In appropriately selected large population screens, highly penetrant genetic variants can be identified easily, as can their impacts on specific aspects of disease outcome. In contrast, targeted mapping crosses between highly discordant strains can help to identify more complex genetic interaction networks such as variants that are penetrant only in the context of specific genetic backgrounds, or epistatic (gene-gene) interaction networks.

We leveraged large-scale population mapping as well as focused intercrosses to better characterize the genetic susceptibility landscape of Sarbecovirus infections in mouse models^26,45^ and demonstrated that a large number of polymorphic loci (**Figure 6**) regulate the host disease responses to this subgroup of coronaviruses. Moreover, in this study and others, we have identified specific genes (*Trim55^26^, Ticam2^45^*, and here *Trim14* and *CCR9*), which have naturally occurring polymorphisms driving aberrant SARS-CoV disease responses. Importantly, we demonstrate that *HrS10* and *HrS11* influence disease severity following both clade I SARS-MA and clade II HKU3-MA infection in the CC-RIX, supporting the hypothesis that intrinsic virulence properties encoded within the Sarbecoviruses are subject to similar susceptibility loci in mammals. Further, the concordant susceptibility profiles of CC011, CC074, *Trim14* and *CCR9* deficient mice with SARS-MA and SARS-CoV-2 MA10 highlight the utility of pre-emergence disease models. Such findings are consistent with the discovery that group I and II human norovirus infection and pathogenesis are heavily regulated by polymorphisms in fucosyltransferase 2 (FUT2)^48^. Rich and complex datasets like the ones described here enable comparisons with human GWAS studies mapping QTL after SARS-CoV-2 infection^33–35^. The CC platform can be used to evaluate the role of these loci in mouse models. One advantage of our approach is the use of different mapping platforms, which provide opportunities to combine datasets across projects to gain a greater understanding of the role of how host genetic variation modulates CoV disease responses in mammals (**Figure 6, Table 1**). In addition, the unexplained heritability and suggestive loci (**Table S1**) we have identified, suggests that CoV disease and immunity are complex polygenic traits, with the accumulation of variants across many loci driving final disease susceptibility. Collectively, these studies represent the most comprehensive analysis of susceptibility loci for an entire genus of human pathogens, identify a large collection of susceptibility loci and candidate genes that regulate multiple aspects type-specific and cross-CoV pathogenesis, validate a role for the CCR9-CCL25 axis in regulating SARS-CoV-2 disease severity and provide a resource for community wide studies.

**Figure 6.**
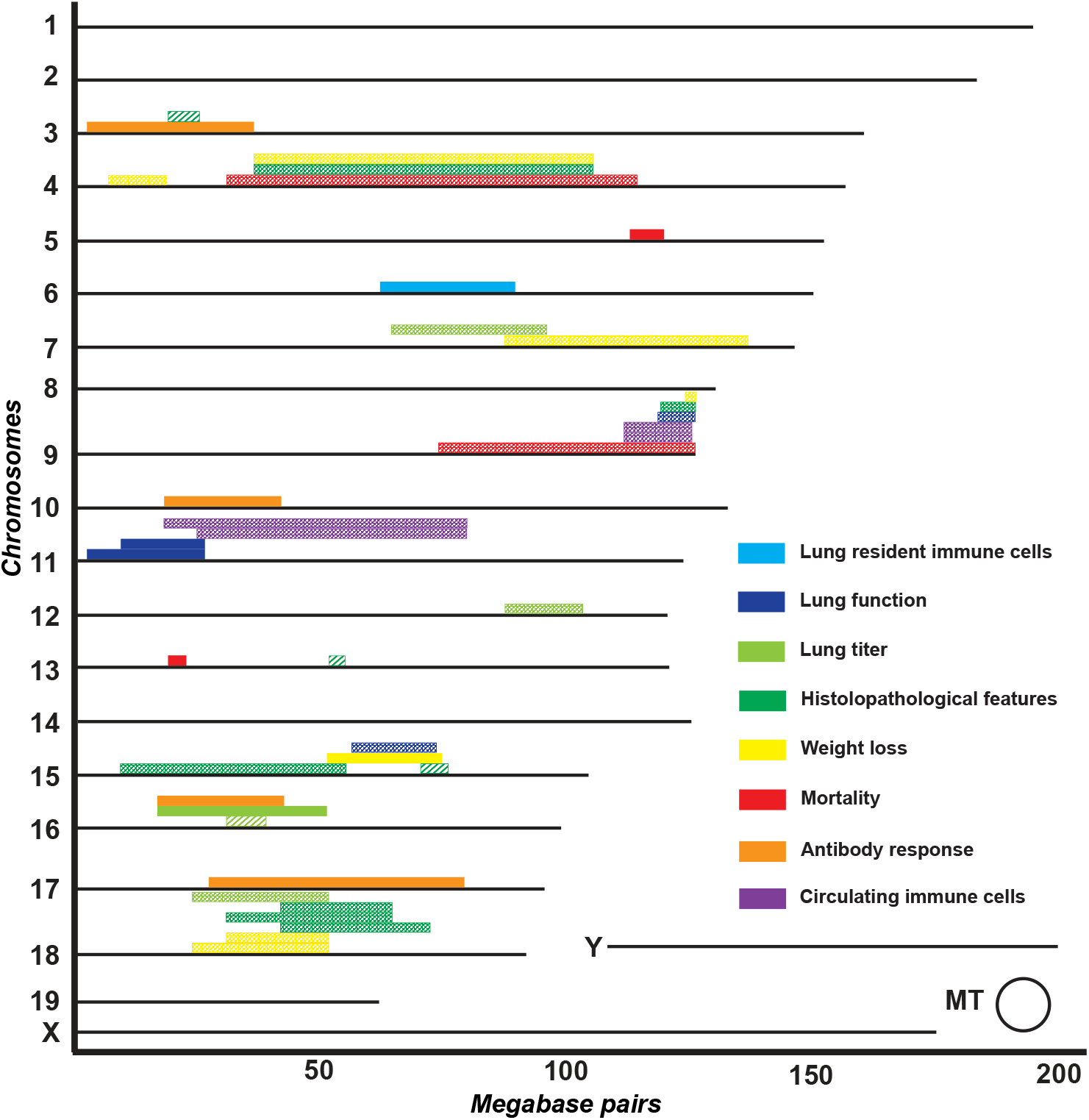
Susceptibility map for SARS-MA and HKU3-MA. Depicted are significant (p=0.95 and p=0.90) QTLs in the Pre-CC (striped), CC-RIX (solid), and the CC011xCC074-F2 and CC003cxCC053-F2 screens, respectively (both checkered).

## Supplemental Figure Legends

**Figure S1.**
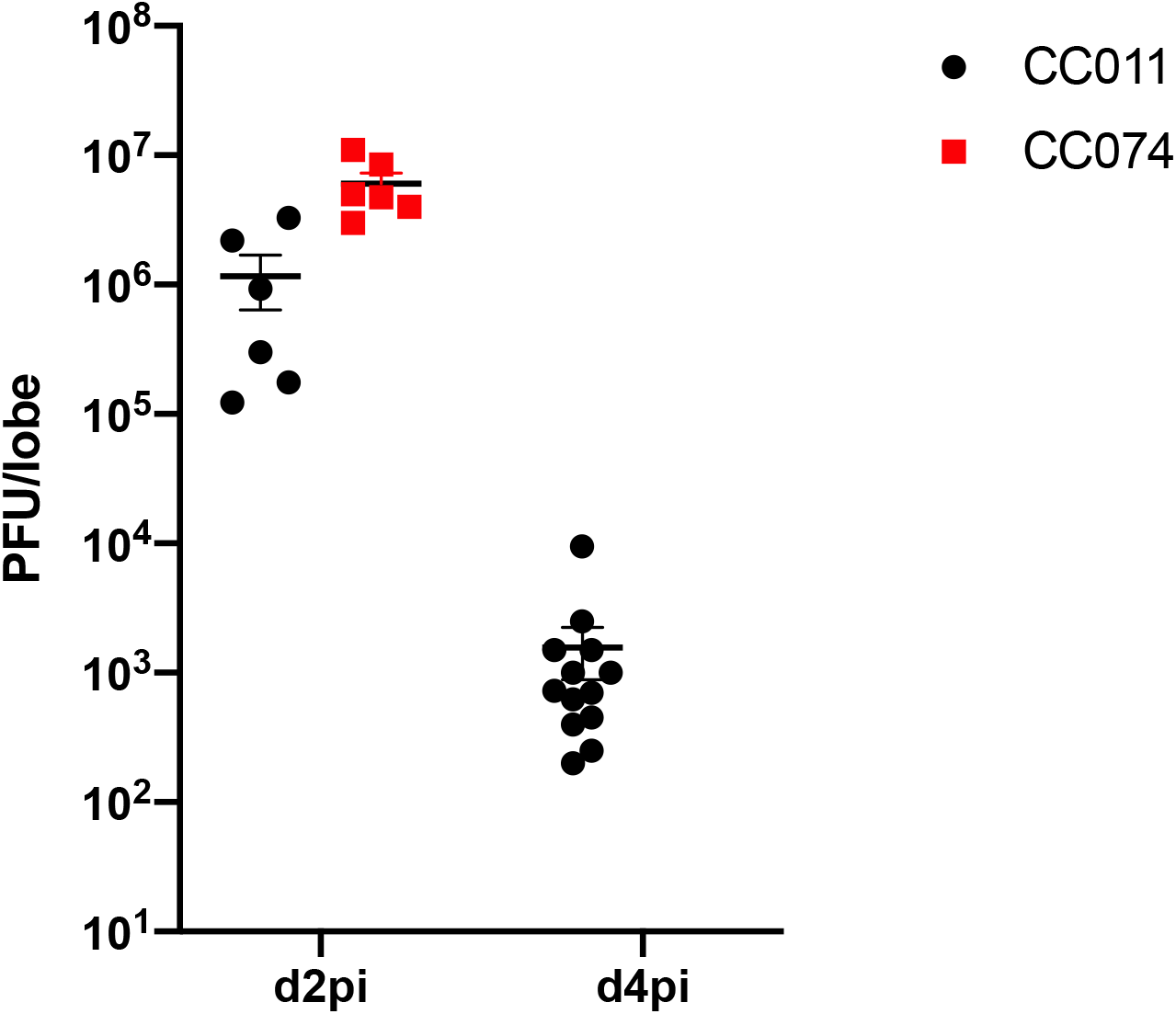
CC074 show elevated viral titer at 2 dpi. Lung tissue of SARS-CoV-2 MA10 infected CC011 (n=6 for d2pi and n=13 for d4pi) and CC074 (n=6 for d2pi and n=13 for d4pi) was tittered by plaque assay to detect viral load on d2pi and d4pi.

**Figure S2.**
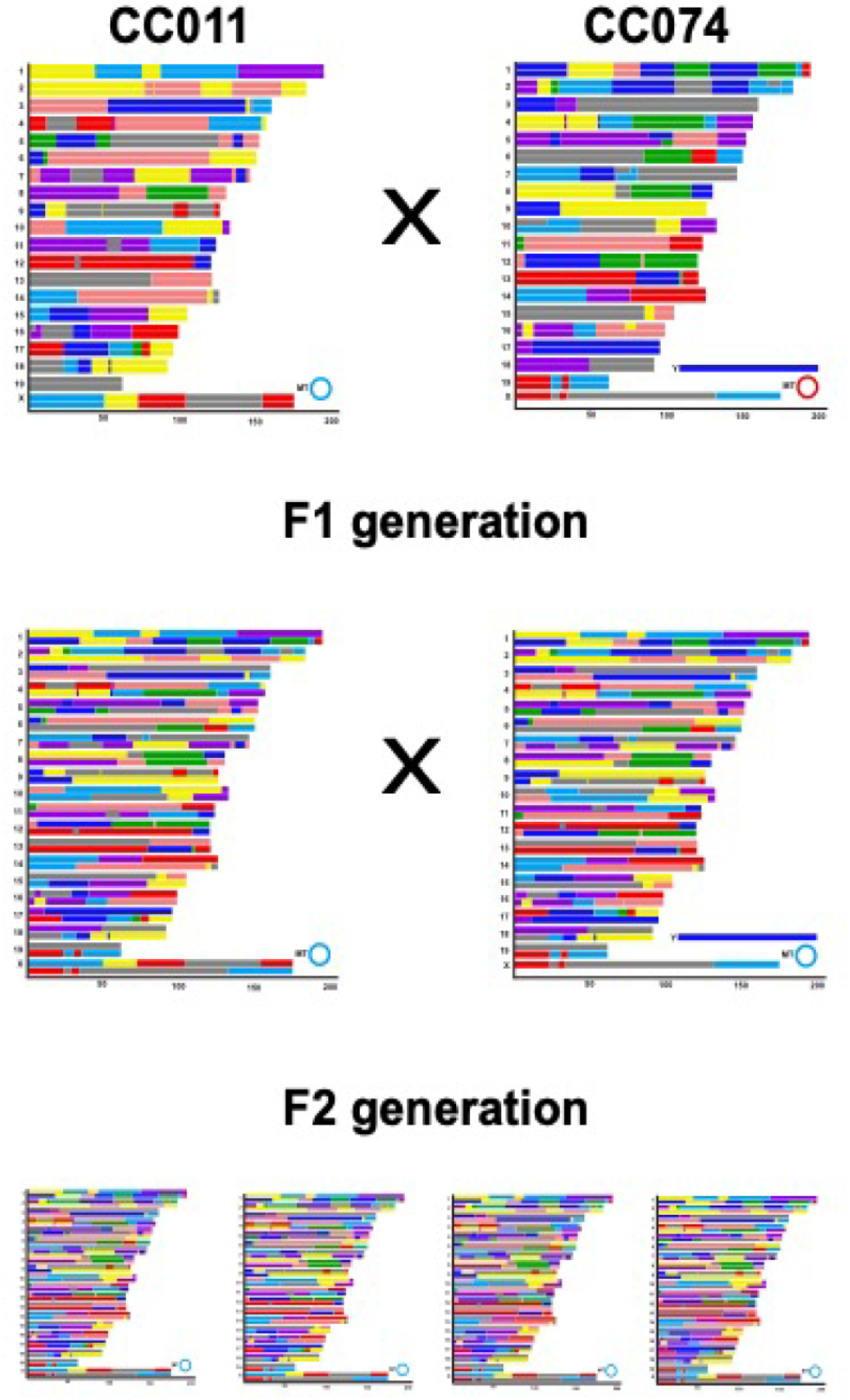
Design of CC011xCC074-F2 cross. In specific cases, individual CC strains may present extreme outlier phenotypes, suggesting a sorting of multiple susceptibility alleles in this strain. In such cases, a mapping intercross (e.g. an F2) between two extreme strains can reveal a large number of causal loci and interaction networks.

**Figure S3.**
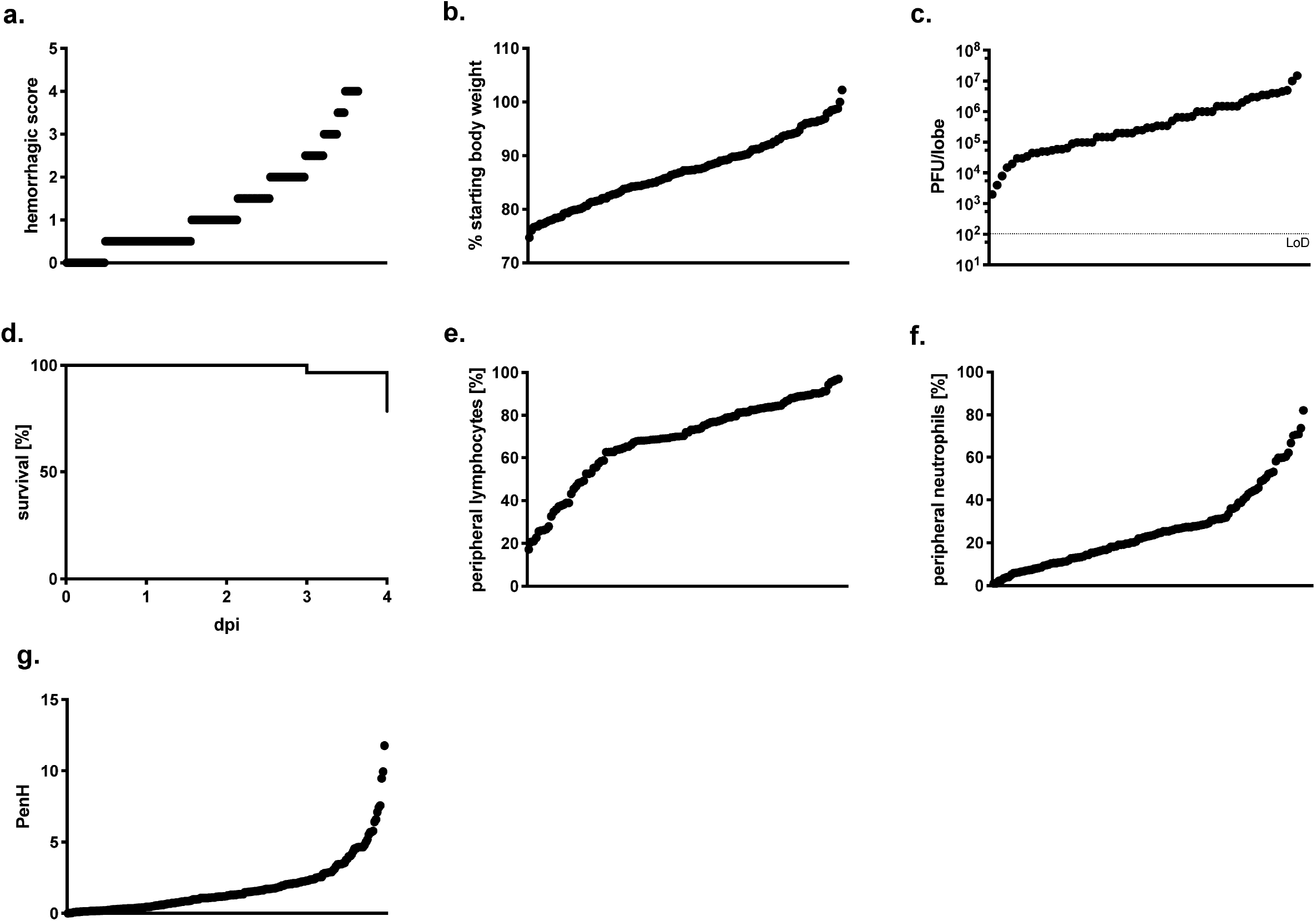
Phenotypic distribution of disease phenotypes after SARS-MA infection in the CC011xCC074-F2 screen. **a.** Lung hemorrhaging, **b.** weight loss at 4 dpi, **c.** lung titer at 4 dpi, **d.** percentage survival, **e.** peripheral blood lymphocytes, **f.** peripheral blood neutrophils, and **g.** PenH lung function. Each dot represents a single animal.

**Figure S4.**
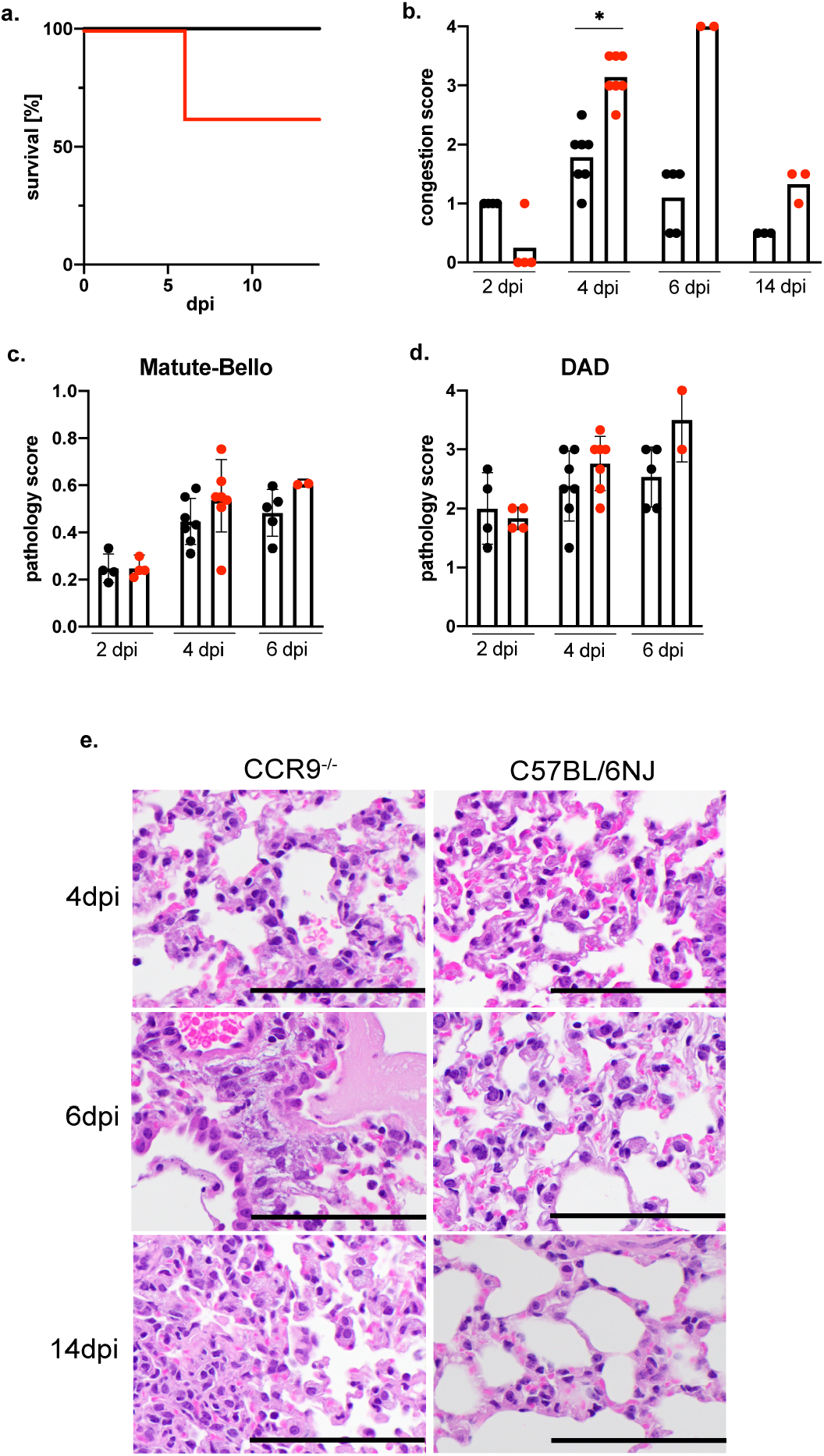
*CCR9^-/-^* mice show significant mortality and lung pathology starting by d6pi of infection. Infection of *CCR9^-/-^* mice with SARS-CoV-2 MA10 showed significant differences in mortality (**a.**), congestion score (**b.**), lung pathology (**c.** and **d.**). Representative H&E stains of lung tissue sections are shown for 4, 6, and 14 dpi for *CCR9^-/-^* and C57BL/6NJ, scale bar indicates 100μm (**e.**), (n=19 CCR9^-/-^ and n=19 C57BL/6NJ; with n=4 for 2 dpi, n=7 for 4 dpi, n=5 for 6 dpi, and n=3 for 14 dpi congestion score and histopathology scoring. Data were analyzed using log-rank (mortality) and Mann-Whitney test (congestion score and pathology scores), *p<0.05.

**Figure S5.**
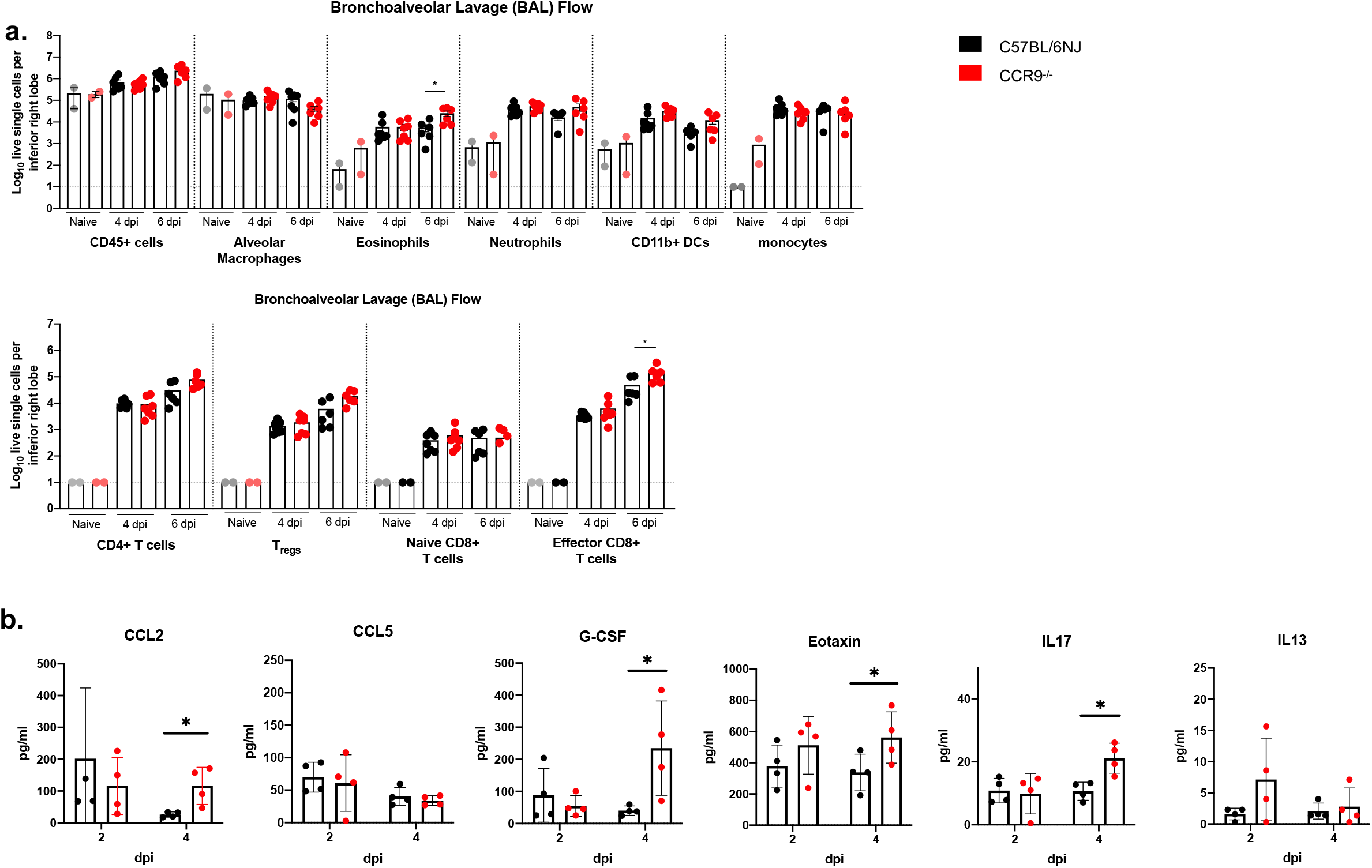
Infiltrating cells in BAL of *CCR9^-/-^* and C57BL/6NJ mice after SARS-CoV-2 MA10 infection. Infection of *CCR9^-/-^* mice with SARS-CoV-2 MA10 showed significant differences in the compositions of lung infiltrating immune cells in BAL (**a.**) as well as in cytokine/chemokine distribution in serum (**b.**) (n=19 CCR9^-/-^ and n=19 C57BL/6NJ; with n=4 for 2 dpi, n=7 for 4 dpi, n=5 for 6 dpi, and n=3 for 14 dpi for multiplex immune-assay and n=12 CCR9^-/-^ and n=13 C57BL/6NJ with n=2 each for mock, n=5-6 for 4 dpi, and n=5 for 6 dpi for analysis of infiltrating cells). Data were analyzed using Mann-Whitney test, *p<0.05, **p<0.01.

**Figure S6.**
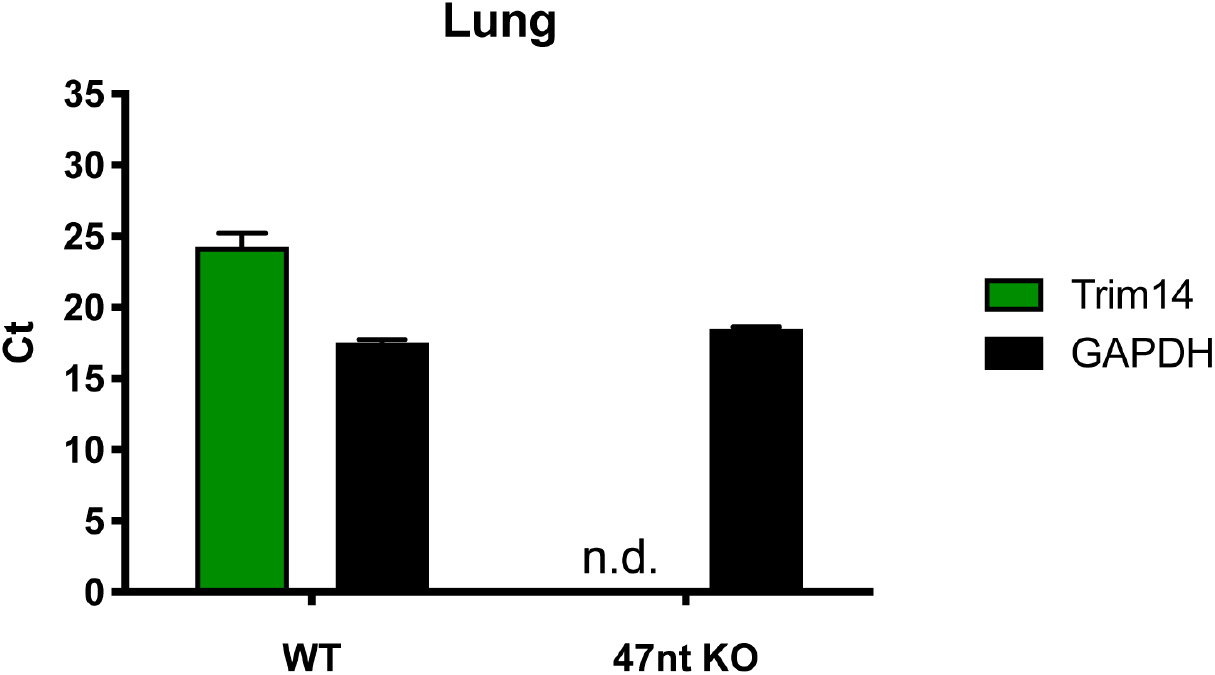
Lung tissue from *Trim14^Δ47/Δ47^* mice lacked detectable Trim14 mRNA. Tissue from *Trim14^Δ47/Δ47^* were found to lack detectable Trim14 mRNA, likely due to nonsense-mediated decay, as measured by RT-qPCR in comparison to littermates.

**Figure S7.**
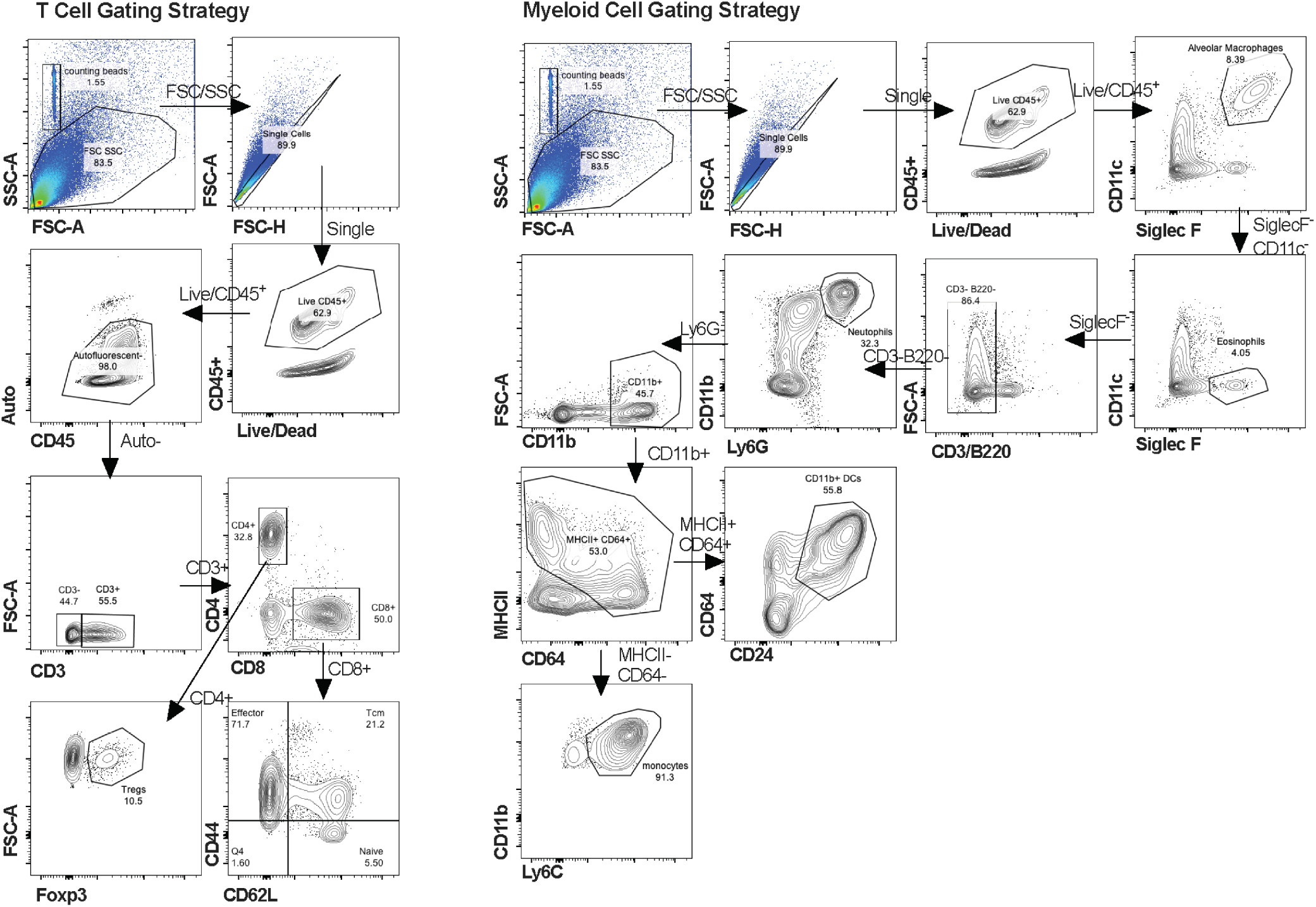
Gating schemes for flow cytometry analysis.

**Supplemental Table S1.** List of suggestive QTLs.

| QTL ID | Chromosome [Mb] | Phenotype(s) (days post infection – dpi) | Haplotype(s)<sup>+</sup> |
| --- | --- | --- | --- |
| **CC011xCC074-F2 suggestive loci** |  |  |  |
| *HrS29* | chr3: 7.2-159.1 | Mortality (4dpi)<br>Hemorrhage (4dpi) | CC011: C, C/D, D, A, D, E<br>CC074: D, H, B, B/F, B/D, B |
| *HrS30* | chr9: 28.2-124.1 | Weight loss (4dpi)<br>Mortality (4dpi)<br>Weight loss females (4dpi) | CC011: B, G, B<br>CC074: A |
| *HrS31* | chr13: 31.9-90.3 | Hemorrhage (4dpi)<br>Periph. monocytes (4dpi) | CC011: B, C<br>CC074: G, D |
| *HrS32* | chr14: 33.8-114.8 | PenH (2dpi) | CC011: C<br>CC074: E, H, G |
| *HrS33* | chrX: 6.2- 93.9 | Rpef (2dpi)<br>Periph. neutrophils (2dpi) | CC011: E, C/E, E, A, G<br>CC074: G, B |
| **CC003xCC053-F2 suggestive loci** |  |  |  |
| *HrS34* | chr3: 58- 148.4 | Weight loss (3dpi) | CC003: B, G, H, G<br>CC053: C/D, C, D, A, E |
| *HrS35* | chr17: 6- 79 | Weight loss (3dpi) | CC003: C/D, C, C/D, D, B<br>CC053: G, E, G, H, A, G, H |
| *HrS36* | chr9: 3.6Mb-74.9Mb | Weight loss (4dpi),<br>Titer (4dpi),<br>Edema (4dpi) | CC003: C, B<br>CC053: B, E |
| *HrS37* | chr10: 5.9Mb-86.3Mb | Titer (4dpi) | CC003: C, H<br>CC053: H, B, A |
| *HrS38* | chrX: 9.4Mb-169.5Mb | Titer (4dpi) | CC003: C, G, C/G, C/E, C, E<br>CC053: A, C, B, F |
| *HrS39* | chr2: 4.3Mb-181.8Mb | mortality (4dpi) | CC003: D, B, E<br>CC053: B, E, G, E, A, C, F |
| *HrS41* | chr6: 34.2Mb-114Mb | Interstitial septum (4dpi),<br>Airspace inflammation (dpi4),<br>Eosinophils (4dpi) | CC003: B, E<br>CC053: G, A, A/D, G |
| *HrS42* | chr11: 37Mb-121.6Mb | Edema (4dpi) | CC003: A/C, B/C, A/C, B/H, B, E, H, C, E<br>CC053: B |
| <sup>+</sup>Haplotype effects are described for each QTL. For F2 crosses, the haplotypes listed are those present in the given parent strains from the proximal to distal ends of each region (A=A/J, B=C57BL/6J, C=129S1/SvImJ, D=NOD/ShiLtJ, E=NZO/HiLtJ, F=CAST/EiJ, G=PWK/PhJ, H=WSB/EiJ). In |  |  |  |

## Acknowledgments

This study was supported by grants in aid from the National Institutes of Health, Allergy and Infectious Diseases (AI100625 and AI149644 to R.S.B., M.T.H., M.T.F., and F.P.M.V.), (R00AG049092 to V.D.M.), a contract from the NIH (HHSN272201700036I; Task Order #38 75N93020F00001 to R.S.B), and support from NCATS (UL1TR002369 to M.A.M. and S.K.M.). We would like to thank the Systems Genetics Core Facility (UNC) for maintaining and distributing Collaborative Cross mice. The research was also supported by a generous gift from the Chan-Zuckerberg foundation.

## Author contributions

A.S., L.E.G., F.P.M.V., S.K.M., M.T.H., V.D.M., M.T.F., R.S.B designed screens and biology experiments; A.S., L.E.G., S.R.L., V.D.M. conducted and characterized infections *in vivo;* A.S., L.E.G., S.R.L., E.S.W., K.L.J., D.T.S. performed *in vitro* studies and immune cell quantifications; A.S., L.E.G, S.R.L., R.L.G., S.A.M., V.D.M., M.T.F. processed and analyzed data, and generated figures; A.S., B.K.H, M.A.M., S.J., S.C., S.K.M., M.T.F. performed genetic mapping studies; L. A.V., L.B.T., M.S.D. designed and isolated *Trim14*-deficient mice; A.S., L.E.G., V.D.M. performed *in vivo* evaluation of *Trim14*-deficient infected animals; S.A.M. scored pathologic changes in the lungs of infected mice; R.L.G., S.A. isolated recombinant viruses; T.A.B., P.H., G.D.S., D.R.M. produced CC-RIX and F2 animals, and processed samples for genotyping; L.E.G., S.R.L., B.K.H, M.A.M., K.L.J., S.J., T.A.B., L.B.T., D.R.M., G.D.S., M.S.D., F.P.M.V., S.K.M., M. T.H. edited the manuscript; A.S., M.T.F., R.S.B. wrote the manuscript.

## Online Material and Methods

### Cells and viruses

Recombinant mouse-adapted SARS-CoV MA15 (SARS-MA), HKU3-SRBD-MA (HKU3-MA), and SARS-CoV-2 MA10 (SARS-2-MA) virus were generated as described previously (HKU3-SRBD-MA: GenBank Accession Number XXX, SARS-CoV-2 MA10: GenBank Accession Number XXX)^11,18,20^. For virus titration, the caudal lobe of the right lung was homogenized in PBS, resulting homogenate was serial-diluted and inoculated onto confluent monolayers of Vero E6 cells (ATCC XXX), followed by agarose overlay. Plaques were visualized with overlay of Neutral Red dye on day 2 (SARS-MA) or day 3 (SARS-CoV-2 MA10) post infection.

### Mouse studies and *in vivo* infections

All mouse studies were performed at the University of North Carolina (Animal Welfare Assurance #A3410-01) using protocols approved by the UNC Institutional Animal Care and Use Committee (IACUC). Animal studies at Washington University were carried out in accordance with the recommendations in the Guide for the Care and Use of Laboratory Animals of the National Institutes of Health. The protocols were approved by the IACUC at the Washington University School of Medicine (Assurance number A3381-01).

Mouse studies fall into three major classes: CC-RIX, F2 intercross mice, and inbred wild-type or gene-edited mice. The laboratory of Pardo Manuel de Villena (FPMV) purchased CC mice from the Systems Genetics Core Facility at UNC between 2012 and 2018. These CC mice were used to breed CC-RIXs in the FPMV laboratory, to ensure proper cohorts and batch sizes. CC-RIXs were generated in a ring design such that each CC-RIX had one copy of the MHC H2B^b^ allele, and that each CC strain was used as both dam and sire in equal proportion across all RIXs. Mice (~105 CC-RIX strains, 3 animals each) were transferred at 5-6 weeks of age to the Baric (RSB) laboratory for infection between 9-12 weeks of age. The details of the CC003 x CC053 F2 are published^45^. The Systems Genetics Core Facility was contracted to generate the F2 cross between CC011 and CC074. F1 mice between CC011 and CC074 were generated in both potential cross directions, and F2 mice were bred in all 4 possible F1 x F1 combinations, to ensure appropriately balanced sex chromosome and parent-of-origin effects. F2 mice (226 males, 177 females) were weaned such that littermates were randomized to different experimental cages to further reduce litter- or batch-effects on the study, and mice were transferred at 5-6 weeks of age to the RSB laboratory for infection between 9-12 weeks of age.

15- week old *CCR9^-/-^* mice (strain 027041) and 15-week old female C57BL/6NJ mice (strain 005304) were purchased from Jackson Laboratory. CC-RIX, CC-F2 mice, *Trim14*-deficient, and CCR9^-/-^ mice were infected with 5×10^3^ (CC-RIX with SARS-MA), 1×10^4^ (CC-F2 with SARS-MA and SARS-CoV-2 MA10), and 1×10^5^ (CC-RIX with HKU3-MA, *Trim14^Δ47/Δ47^* and *CCR9^-/-^* mice with SARS-MA and SARS-CoV-2 MA10) plaque forming units (PFU) in 50 μl PBS intranasally at 9-12 (CC-RIX and CC-F2 mice) or 15 (*CCR9^-/-^* and C57BL/6NJ) weeks of age, respectively. Body weight, mortality, and pulmonary function by whole body plethysmography^49^ were monitored daily where indicated. At indicated timepoints, mice were euthanized and gross pathology (hemorrhage score) of the lung was assessed and scored on a scale from 0 (no hemorrhage) to 4 (severe hemorrhage affecting all lung lobes). Then lung tissue was harvested for titer and histopathology analysis; and blood samples were harvested to determine antibody composition and for analysis of peripheral immune cells. Samples were stored at −80°C until homogenized and titered by plaque assay as described above. Serum was prepared and SARS-CoV spike-specific antibody were quantified by ELISA as previously described^33^. Peripheral blood was diluted 1:5 in PBS/EDTA and analyzed with the VetScan HM5 as previously described ^50^. Histopathology samples were fixed in 10% phosphate buffered formalin for 7 days before paraffin embedding, sectioning stained with hematoxylin and eosin.

### Generation of *Trim14-deficient* mice

Gene-edited *Trim14*-deficient mice were generated with support from the Genome Engineering and iPSC center and Department of Pathology Micro-Injection Core (Washington University School of Medicine). A sgRNA targeting exon 4 of *Trim14* was selected based on minimal off-target effects *in silico* and targeting efficiency *in vitro*. The sgRNA (5’-ACCAATGGACACTCGCCTGANGG-3’) was synthesized, transcribed (HiScribe T7 *In vitro* Transcription Kit, New England BioLabs), and purified (MEGAclear Transcription Clean-Up Kit, Thermo Fisher). The sgRNA was mixed and co-injected with Cas9 RNA at 5ng/μl and 10ng/μl final concentrations into half-day-old C57BL/6J embryos (E0.5). After next-generation sequencing of founders and two generations of mice backcrossed to C57BL/6J mice, a mouse line with a 47-nucleotide deletion (5’-GCCTGAAGGAAAGTGAGTTGCCTAAGACCAACTCCAAGTCCTTGCTC-3’) encompassing the 3’ splice site of intron 4 and part of the coding region of exon 4 was generated. These *Trim14^Δ47/Δ47^* mice were bred as homozygotes and used for experiments. *Trim14^Δ47/Δ47^* mice were born in normal Mendelian frequencies and showed no apparent defects in development, growth, or fecundity. Lung tissue from *Trim14^Δ47/Δ47^* were found to lack detectable *Trim14* mRNA, likely due to nonsense-mediated decay, as measured by RT-qPCR using a predesigned primer/probe set for *Trim14* (IDT, Assay ID Mm.PT.58.286730) and the housekeeping gene *GAPDH* (IDT, Assay ID Mm.PT.39a.1). Sanger sequencing of a polymerase chain reaction amplicon [5’ GGCACAGCTCAACCCATGG −3’ (forward) and 5’-ACCAGCGAGCTCGTGCTCC −3’ (reverse)] was used for genotyping.

### Flow cytometry analysis of immune cell infiltrates

For analysis of BAL fluid, mice were sacrificed by ketamine overdose, followed by cannulation of the trachea with a 19-G canula. BAL was performed with three washes of 0.8 ml of sterile PBS. BAL fluid was centrifuged, and single cell suspensions were generated for staining. For analysis of lung tissues, mice were perfused with sterile PBS, and the right inferior lung lobes were digested at 37°C with 630 μg/mL collagenase D (Roche) and 75 U/mL of DNase I (Sigma–Aldrich) for 2 h. Single cell suspensions of BAL fluid and lung digest were preincubated with Fc Block antibody (BD PharMingen) in PBS + 2% heat-inactivated FBS for 10 min at room temperature before staining. Cells were incubated with antibodies against the following markers: efluor506 Viability Dye (Thermo Fisher, 65-0866-14), BUV395 anti-CD45 (Clone 30-F11, BD Biosciences), BV711 anti-CD11b (Clone M1/70, Biolegend), APC-Cy7 anti-CD11c (Clone HL3, BD Biosciences), BV650 anti-Ly6G (Clone 1A8, Biolegend), Pacific Blue anti-Ly6C (Clone HK1.4, Biolegend) FITC anti-CD24 (Clone M1/69, Biolegend), PE anti-Siglec F (Clone E50-2440, Biolegend), PE-Cy7 anti-CD64 (Clone X54-5/7.1, Biolegend), AF700 anti-MHCII (Clone M5/114.15.2, Biolegend), BV421 anti-CD3 (Clone 17A2, Biolegend), BV785 anti-CD4 (Clone GK1.5, Biolegend), APC anti-CD8a (Clone 53-6.7, Biolegend) BV421 anti-B220 (Clone RA3-6B2, Biolegend) APC-Cy7 anti-CD44 (Clone IM7, Biolegend) BV605 anti-CD62L (Clone MEL-14, Biolegend). All antibodies were used at a dilution of 1:200. Cells were stained for 20 min at 4°C, washed, fixed and permeabilized for intracellular staining with Foxp3/Transcription Factor Staining Buffer Set (eBioscience) according to manufacturer’s instructions. Cells were incubated overnight at 4°C with BV421 anti-Foxp3 (Clone MF-14, Biolegend) washed, re-fixed with 4% PFA (EMS) for 20 min and resuspended in permeabilization buffer. Absolute cell counts were determined using Trucount beads (BD). Flow cytometric data were acquired on a cytometer (BD-X20; BD Biosciences) and analyzed using FlowJo software (Tree Star) (**Figure S7**).

### Cytokine and chemokine protein analysis

The small center lung lobe of each mouse was homogenized in 1 ml of PBS and briefly centrifuged to remove debris. Fifty microliters of homogenate were used to measure cytokine and chemokine protein abundance using a Bio-Plex Pro mouse cytokine 23-plex assay (Bio-Rad) according to the manufacturer’s instructions.

### Lung pathology scoring

Two separate lung pathology scoring scales, Matute-Bello and Diffuse Alveolar Damage (DAD), were used to quantify acute lung injury (ALI)^51^.

For Matute-Bello scoring samples were blinded and three random fields of lung tissue were chosen and scored for the following: (A) neutrophils in alveolar space (none = 0, 1–5 cells = 1, > 5 cells = 2), (B) neutrophils in interstitial space (none = 0, 1–5 cells = 1, > 5 cells = 2), (C) hyaline membranes (none = 0, one membrane = 1, > 1 membrane = 2), (D) Proteinaceous debris in air spaces (none = 0, one instance = 1, > 1 instance = 2), (E) alveolar septal thickening (< 2Å~ mock thickness = 0, 2–4Å~ mock thickness = 1, > 4Å~ mock thickness = 2). Scores from A–E were put into the following formula score = [(20x A) + (14 x B) + (7 x C) + (7 x D) + (2 x E)]/100 to obtain a lung injury score per field and then averaged for the final score for that sample.

In a similar way, for DAD scoring, three random fields of lung tissue were scored for the in a blinded manner for: 1= absence of cellular sloughing and necrosis, 2= uncommon solitary cell sloughing and necrosis (1–2 foci/field), 3=multifocal (3+foci) cellular sloughing and necrosis with uncommon septal wall hyalinization, or 4=multifocal (>75% of field) cellular sloughing and necrosis with common and/or prominent hyaline membranes. To obtain the final DAD score per mouse, the scores for the three fields per mouse were averaged.

### Genotyping

CC003, CC053, their F1 progeny, and the F2 cross were genotyped as previously described^45^. CC011, CC074, their F1 progeny, and the F2 cross were genotyped on the MiniMUGA genotyping array^52^. Genomic DNA was isolated from tail-clips of animals using the Qiagen (Hilden, Germany) DNeasy Blood & Tissue kit. 1.5 μg was sent to Neogen (Lincoln, Nebraska) for processing. We filtered the genotypes upon return for informativeness within this cross. To be considered informative, the marker had to have one homozygous allele in all CC011 mice genotyped, the alternate homozygous allele in all CC074 mice genotyped, and the appropriate call in all F1 animals (H calls on the autosomes, an H call in females on the X chromosomes, and the relevant homozygous call in male F1s). This filtering reduced the ~10,800 SNPs on the MiniMUGA array to 2821 informative markers.

### QTL mapping and statistical analyses

For the CC-RIX, we used the same pipeline we previously described^29^. Briefly, each CC-RIX had their genome represented as an array of probabilities of each of the 8 CC founder haplotypes (**Figure 1C**). This array was used in the DOQTL R package^53^ to run an 8-allele regression at each of 77,000 markers for our CC-RIX phenotypes. At each marker, a LOD score is calculated describing the goodness of fit of our trait~genotype model relative to a null model. Significance was determined by running 1000 permutations scrambling the relationship between phenotypes and haplotypes. In this way, significance is independent of both population allele frequencies, as well as the phenotypic distribution. For the F2 crosses, instead of a regression on haplotype probabilities, the R/QTL package conducts a regression of the trait of interest on the exact genotypes at each locus^54^. As with the CC-RIX mapping, permutation testing is used to identify significance.

## References

1 Ge, D. et al. Genetic variation in IL28B predicts hepatitis C treatment-induced viral clearance. Nature 461, 399–401, doi:10.1038/nature08309 (2009).

2 McLaren, P. J. et al. Polymorphisms of large effect explain the majority of the host genetic contribution to variation of HIV-1 virus load. Proc Natl Acad Sci U S A 112, 14658–14663, doi:10.1073/pnas.1514867112 (2015).

3 Wang, Q., Vlasova, A. N., Kenney, S. P. & Saif, L. J. Emerging and re-emerging coronaviruses in pigs. Curr Opin Virol 34, 39–49, doi:10.1016/j.coviro.2018.12.001 (2019).

4 Chen, B. et al. Overview of lethal human coronaviruses. Signal Transduct Target Ther 5, 89, doi:10.1038/s41392-020-0190-2 (2020).

5 Zhou, P. et al. A pneumonia outbreak associated with a new coronavirus of probable bat origin. Nature 579, 270–273, doi:10.1038/s41586-020-2012-7 (2020).

6 Zhang, Y. Z. & Holmes, E. C. A Genomic Perspective on the Origin and Emergence of SARS-CoV-2. Cell 181, 223–227, doi:10.1016/j.cell.2020.03.035 (2020).

7 Menachery, V. D. et al. SARS-like WIV1-CoV poised for human emergence. Proc Natl Acad Sci U S A 113, 3048–3053, doi:10.1073/pnas.1517719113 (2016).

8 Menachery, V. D. et al. A SARS-like cluster of circulating bat coronaviruses shows potential for human emergence. Nat Med 21, 1508–1513, doi:10.1038/nm.3985 (2015).

9 Anthony, S. J. et al. Further Evidence for Bats as the Evolutionary Source of Middle East Respiratory Syndrome Coronavirus. mBio 8, doi:10.1128/mBio.00373-17 (2017).

10 Coronaviridae Study Group of the International Committee on Taxonomy of, V. The species Severe acute respiratory syndrome-related coronavirus: classifying 2019-nCoV and naming it SARS-CoV-2. Nat Microbiol 5, 536–544, doi:10.1038/s41564-020-0695-z (2020).

11 Becker, M. M. et al. Synthetic recombinant bat SARS-like coronavirus is infectious in cultured cells and in mice. Proc Natl Acad Sci U S A 105, 19944–19949, doi:10.1073/pnas.0808116105 (2008).

12 Ahmad, T., Haroon, Baig, M. & Hui, J. Coronavirus Disease 2019 (COVID-19) Pandemic and Economic Impact. Pak J Med Sci 36, S73–S78, doi:10.12669/pjms.36.COVID19-S4.2638 (2020).

13 Enserink, M. & Kupferschmidt, K. With COVID-19, modeling takes on life and death importance. Science 367, 1414–1415, doi:10.1126/science.367.6485.1414-b (2020).

14 Rasmussen, A. L. et al. Host genetic diversity enables Ebola hemorrhagic fever pathogenesis and resistance. Science 346, 987–991, doi:10.1126/science.1259595 (2014).

15 Sanchez, A., Wagoner, K. E. & Rollin, P. E. Sequence-based human leukocyte antigen-B typing of patients infected with Ebola virus in Uganda in 2000: identification of alleles associated with fatal and nonfatal disease outcomes. J Infect Dis 196 Suppl 2, S329–336, doi:10.1086/520588 (2007).

16 Cameron, M. J. et al. Interferon-mediated immunopathological events are associated with atypical innate and adaptive immune responses in patients with severe acute respiratory syndrome. J Virol 81, 8692–8706, doi:10.1128/JVI.00527-07 (2007).

17 Shang, J. et al. Structure of mouse coronavirus spike protein complexed with receptor reveals mechanism for viral entry. PLoS Pathog 16, e1008392, doi:10.1371/journal.ppat.1008392 (2020).

18 Roberts, A. et al. A mouse-adapted SARS-coronavirus causes disease and mortality in BALB/c mice. PLoS Pathog 3, e5, doi:10.1371/journal.ppat.0030005 (2007).

19 Dinnon, K. H. et al. A mouse-adapted SARS-CoV-2 model for the evaluation of COVID-19 medical countermeasures. bioRxiv, doi:10.1101/2020.05.06.081497 (2020).

20 Leist, S. R. et al. A Mouse-adapted SARS-CoV-2 induces Acute Lung Injury (ALI) and mortality in Standard Laboratory Mice. Cell, doi:10.1016/j.cell.2020.09.050 (2020).

21 Leist, S. R. & Baric, R. S. Giving the Genes a Shuffle: Using Natural Variation to Understand Host Genetic Contributions to Viral Infections. Trends Genet 34, 777–789, doi:10.1016/j.tig.2018.07.005 (2018).

22 Schafer, A., Baric, R. S. & Ferris, M. T. Systems approaches to Coronavirus pathogenesis. Curr Opin Virol 6, 61–69, doi:10.1016/j.coviro.2014.04.007 (2014).

23 Noll, K. E., Ferris, M. T. & Heise, M. T. The Collaborative Cross: A Systems Genetics Resource for Studying Host-Pathogen Interactions. Cell Host Microbe 25, 484–498, doi:10.1016/j.chom.2019.03.009 (2019).

24 Srivastava, A. et al. Genomes of the Mouse Collaborative Cross. Genetics 206, 537–556, doi:10.1534/genetics.116.198838 (2017).

25 Collaborative Cross, C. The genome architecture of the Collaborative Cross mouse genetic reference population. Genetics 190, 389–401, doi:10.1534/genetics.111.132639 (2012).

26 Gralinski, L. E. et al. Genome Wide Identification of SARS-CoV Susceptibility Loci Using the Collaborative Cross. PLoS Genet 11, e1005504, doi:10.1371/journal.pgen.1005504 (2015).

27 Frieman, M. et al. Molecular determinants of severe acute respiratory syndrome coronavirus pathogenesis and virulence in young and aged mouse models of human disease. J Virol 86, 884–897, doi:10.1128/JVI.05957-11 (2012).

28 Maurizio, P. L. et al. Bayesian Diallel Analysis Reveals Mx1-Dependent and Mx1-Independent Effects on Response to Influenza A Virus in Mice. G3 (Bethesda) 8, 427–445, doi:10.1534/g3.117.300438 (2018).

29 Noll, K. E. et al. Complex Genetic Architecture Underlies Regulation of Influenza-A-Virus-Specific Antibody Responses in the Collaborative Cross. Cell Rep 31, 107587, doi:10.1016/j.celrep.2020.107587 (2020).

30 Aylor, D. L. et al. Genetic analysis of complex traits in the emerging Collaborative Cross. Genome Res 21, 1213–1222, doi:10.1101/gr.111310.110 (2011).

31 Ferris, M. T. et al. Modeling host genetic regulation of influenza pathogenesis in the collaborative cross. PLoS Pathog 9, e1003196, doi:10.1371/journal.ppat.1003196 (2013).

32 Abiola, O. et al. The nature and identification of quantitative trait loci: a community’s view. Nat Rev Genet 4, 911–916, doi:10.1038/nrg1206 (2003).

33 Ellinghaus, D. et al. Genomewide Association Study of Severe Covid-19 with Respiratory Failure. N Engl J Med, doi:10.1056/NEJMoa2020283 (2020).

34 Pairo-Castineira, E. et al. Genetic mechanisms of critical illness in Covid-19. Nature, doi: 10.1038/s41586-020-03065-y (2020).

35 Shelton, J. F. et al. Trans-ancestry analysis reveals genetic and nongenetic associations with COVID-19 susceptibility and severity. Nat Genet, doi:10.1038/s41588-021-00854-7 (2021).

36 Pathak, M. & Lal, G. The Regulatory Function of CCR9(+) Dendritic Cells in Inflammation and Autoimmunity. Front Immunol 11, 536326, doi:10.3389/fimmu.2020.536326 (2020).

37 Wang, C. et al. The role of chemokine receptor 9/chemokine ligand 25 signaling: From immune cells to cancer cells. Oncol Lett 16, 2071–2077, doi:10.3892/ol.2018.8896 (2018).

38 Lopez-Pacheco, C., Soldevila, G., Du Pont, G., Hernandez-Pando, R. & Garcia-Zepeda, E. A. CCR9 Is a Key Regulator of Early Phases of Allergic Airway Inflammation. Mediators Inflamm 2016, 3635809, doi:10.1155/2016/3635809 (2016).

39 Nakamoto, N. Role of inflammatory macrophages and CCR9/CCL25 chemokine axis in the pathogenesis of liver injury as a therapeutic target. Nihon Rinsho Meneki Gakkai Kaishi 39, 460–467, doi:10.2177/jsci.39.460 (2016).

40 Chu, P. S. et al. C-C motif chemokine receptor 9 positive macrophages activate hepatic stellate cells and promote liver fibrosis in mice. Hepatology 58, 337–350, doi:10.1002/hep.26351 (2013).

41 Channappanavar, R. et al. Dysregulated Type I Interferon and Inflammatory Monocyte-Macrophage Responses Cause Lethal Pneumonia in SARS-CoV-Infected Mice. Cell Host Microbe 19, 181–193, doi:10.1016/j.chom.2016.01.007 (2016).

42 Murdoch, B. M. et al. Genome-wide scan identifies loci associated with classical BSE occurrence. PLoS One 6, e26819, doi:10.1371/journal.pone.0026819 (2011).

43 Prisco, S. Z. et al. Refined mapping of a hypertension susceptibility locus on rat chromosome 12. Hypertension 64, 883–890, doi:10.1161/HYPERTENSIONAHA.114.03550 (2014).

44 Jamieson, S. E. et al. Evidence for a cluster of genes on chromosome 17q11-q21 controlling susceptibility to tuberculosis and leprosy in Brazilians. Genes Immun 5, 46–57, doi:10.1038/sj.gene.6364029 (2004).

45 Gralinski, L. E. et al. Allelic Variation in the Toll-Like Receptor Adaptor Protein Ticam2 Contributes to SARS-Coronavirus Pathogenesis in Mice. G3 (Bethesda) 7, 1653–1663, doi:10.1534/g3.117.041434 (2017).

46 Zhou, Z. et al. TRIM14 is a mitochondrial adaptor that facilitates retinoic acid-inducible gene-I-like receptor-mediated innate immune response. Proc Natl Acad Sci U S A 111, E245–254, doi:10.1073/pnas.1316941111 (2014).

47 Tan, P. et al. Assembly of the WHIP-TRIM14-PPP6C Mitochondrial Complex Promotes RIG-I-Mediated Antiviral Signaling. Mol Cell 68, 293–307 e295, doi:10.1016/j.molcel.2017.09.035 (2017).

48 Lindesmith, L. et al. Human susceptibility and resistance to Norwalk virus infection. Nat Med 9, 548–553, doi:10.1038/nm860 (2003).

49 Menachery, V. D., Gralinski, L. E., Baric, R. S. & Ferris, M. T. New Metrics for Evaluating Viral Respiratory Pathogenesis. PLoS One 10, e0131451, doi:10.1371/journal.pone.0131451 (2015).

50 Leist, S. R., Jensen, K. L., Baric, R. S. & Sheahan, T. P. Increasing the translation of mouse models of MERS coronavirus pathogenesis through kinetic hematological analysis. PLoS One 14, e0220126, doi:10.1371/journal.pone.0220126 (2019).

51 Sheahan, T. P. et al. Comparative therapeutic efficacy of remdesivir and combination lopinavir, ritonavir, and interferon beta against MERS-CoV. Nat Commun 11, 222, doi:10.1038/s41467-019-13940-6 (2020).

52 Sigmon, J. S. et al. Content and performance of the MiniMUGA genotyping array, a new tool to improve rigor and reproducibility in mouse research. bioRxiv, 2020.2003.2012.989400, doi:10.1101/2020.03.12.989400 (2020).

53 Gatti, D. M. et al. Quantitative trait locus mapping methods for diversity outbred mice. G3 (Bethesda) 4, 1623–1633, doi:10.1534/g3.114.013748 (2014).

54 Broman, K. W., Wu, H., Sen, S. & Churchill, G. A. R/qtl: QTL mapping in experimental crosses. Bioinformatics 19, 889–890, doi:10.1093/bioinformatics/btg112 (2003).

